# B Cell-Induced Lymph Node Stromal Remodeling Compromised Neutrophil Response to Secondary *Staphylococcus aureus* Infection

**DOI:** 10.1101/2025.10.31.681727

**Authors:** Jingna Xue, Darellynn Oo, Nicole Rosin, Keerthana Chockalingam, Jeff Biernaskie, Peng Huang, Shan Liao

## Abstract

The lymph node (LN) stromal cells create niches, regulate lymph drainage, and guide the migration, positioning, and activation of immune cells within the LN. Upon stimulation, the stromal cells respond dynamically to accommodate the massive expansion of immune cells in the LN. However, there is still a limited understanding of how niche-associated stromal cell subsets are remodeled and how this remodeling reshapes host immune protection. In an oxazolone (OX)-induced skin inflammation model, we found that LN remodeling weakened the neutrophil response to secondary *Staphylococcus aureus* (*S. aureus*) infection in the inflamed LN, whereas depleting B cells rescued the responses. To understand the mechanism, we used single-cell RNA sequencing (scRNAseq) to characterize alterations in fibroblastic reticular cell (FRC) subsets and identified a new FRC subset expressing an intermediate level of CXCL13 (Cxcl13^int^ RCs) in the inflamed LN. Further studies revealed that Cxcl13^int^ RCs replaced Ccl19^lo^ FRCs in the interfollicular zones (IFZs) of the inflamed LN, leading to a compromised conduit network. Depleting B cells preserved the integrity and function of the FRC-conduit network within the IFZ. Finally, the induction of Cxcl13^int^ RCs depended on the B cell-derived lymphotoxin signaling. The induced Cxcl13^int^ RCs were transitional, creating a time window of comprised immune protection during the OX-inflammation. This study elucidated a mechanism underlying LN stromal remodeling and provides proof-of-principle evidence that B cell-stromal interactions may be a novel target to enhance host immunity in diseased conditions.

## Introduction

*Staphylococcus aureus* (***S. aureus***) is an opportunistic pathogen and one of the most common causes of skin and soft tissue infections (**SSTIs**) in humans. Patients with atopic dermatitis and other skin diseases are associated with a higher risk of *S. aureus* infection and bacteremia^1–4^. The lymph nodes (LNs), serving as a central hub of immune protection, are crucial physical barriers preventing bacterial dissemination^5,6^. Stromal cells in LNs, including endothelial cells and non-endothelial fibroblastic reticular cells (FRCs), form unique niches that support optimal immune responses. Lymphatic endothelial cells (LECs) create the subcapsular sinus (SCS) and medullary sinus (MS), the major niches that guide lymph and immune cell entry into the LN from the afferent lymphatic vessels and exit via the efferent lymphatic vessels. LECs line the subcapsular sinus (SCS) and the medullary sinus (MS), whereas FRCs produce chemokines and cytokines that facilitate lymphocyte migration, positioning, survival, and activation within the LN. Both LEC and FRC populations are highly heterogeneous, with each subset expressing a unique gene signature and closely associated with distinct niches within the LN^7,8,17,9–16^.

Additionally, FRCs construct and maintain a conduit network that directs lymph drainage within the LN by producing and depositing various extracellular matrix (**ECM**) components^7–9^. Under homeostatic conditions, the conduit network guides lymph distribution within the LN, enabling low-molecule-weight materials, including lymph-borne antigens, cytokines, and chemokines, to access cells and high endothelial venules (**HEVs**) in the interfollicular zone (IFZs), T-B cell border, and T cell zone (TCZs) ^7,18–21^. Antigens drained via the conduit network can be directly sampled by LN residential DCs and B cells^19,22,23^. The conduit network also deposits the chemokines and cytokines produced locally or drained from the peripheral tissue to direct immune cell infiltration and migration in the LN^20,21,24^. This step is essential for the first wave of neutrophil infiltration to the LN following *S. aureus* infection^5,24^.

Upon immune stimulation, LNs rapidly enlarge due to the recruitment and proliferation of immune cells. FRCs respond dynamically to adapt to these changes in the LNs. In the acute phase, CLEC-2^+^ dendritic cells (**DCs**) infiltrate the LNs and interact with podoplanin (**PDPN**) expressed by FRCs^25,26^, which rapidly increase their surface area and accommodate the initial phase of LN expansion^25^. Over time, FRCs undergo proliferation to support continuously increasing cell numbers^8,9^. The factors inducing FRC expansion include mechanical stress^27^ and engagement of inflammatory signaling pathways^28–30^, which vary from different stimulations. Global transcriptomic changes in FRC subpopulations following immune stimulations have also been shown by scRNAseq^31,32^. These studies showed that immune stimulation significantly alters the cytokine and chemokine expression profiles in FRCs subpopulations, substantially impacting adaptive immunity. Although multiple studies have shown that inflammation can disrupt the conduit network ^7,22,24,33^, the mechanism by which FRC remodeling compromises conduit integrity remains unclear.

In previous study, we have observed that OX-induced skin inflammation (OX-inflammation) compromised neutrophil response to the secondary *S. aureus* infection and increased the risk of bacterial systemic dissemination. In this study, we found that depleting B cells rescue the neutrophil responses to *S. aureus* within the LN. To understand the mechanism, we performed scRNAseq analysis to identify and subsequently map the remodeling of LEC and FRC subpopulations within the LN using immunofluorescent imaging. Alongside a dramatic change in the gene expression profiles of LEC and FRC subpopulations, we identified an induced FRC subset, expressing an intermediate level of CXCL13 (Cxcl13^int^ RCs), which replaced Ccl19^lo^ FRCs in the IFZ, contributing to the destruction of the conduit network in the inflamed LNs. Depleting B cells preserved the FRC-conduit network within the IFZ. Finally, our results indicated that the emergence of Cxcl13^int^ RCs was driven by B cells and dependent on lymphotoxin signaling.

## Results

### B cell depletion restores the neutrophil response to secondary *S. aureus* infection during OX-inflammation

LNs are highly organized into various niches typically based on the T cell and B cell compartmentalization (**Figure 1A**), and each niche is supported by a unique subset of FRCs (**Figure 1B**). Upon *S. aureus* infection, the prompt recruitment of neutrophils from the HEVs to the LN (<4 hours post-infection, 4hpi) prevents bacterial dissemination into the systemic circulation^5,24^. Our previous study showed that the neutrophil response to *S. aureus* infection was impaired in the inflamed LN at day 4 post OX-induced skin inflammation (OXd4), which corresponds to the peak of OX-inflammation, posing a significant risk of bacterial systemic dissemination^24^.

**Figure 1.**
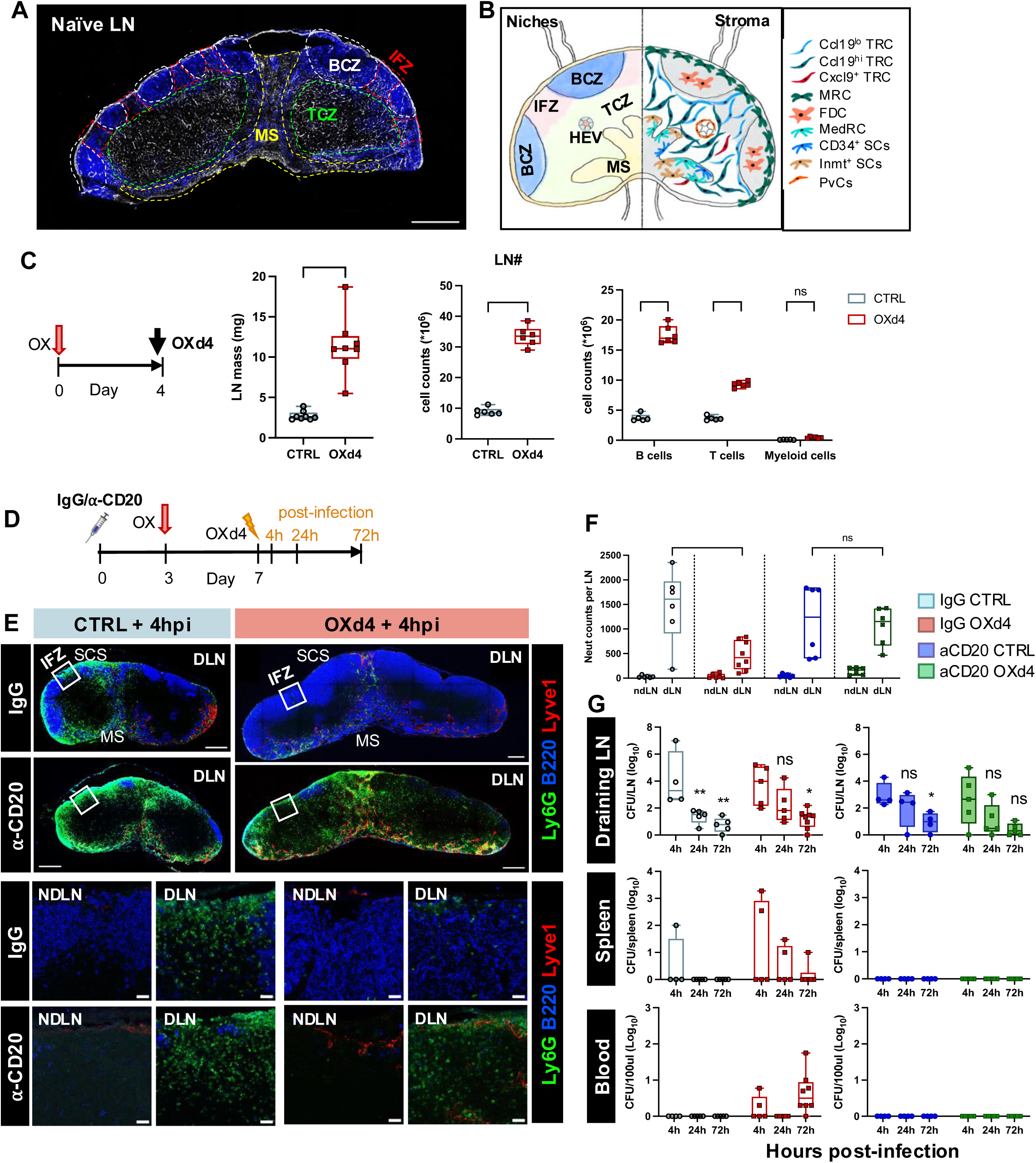
B cell depletion rescued neutrophil response to *S. aureus* during OX-inflammation. (**A**) Representative tile scan image of naïve LN with anti-collagen I (conduit) and anti-B220 (B cells) staining showing LN niches. (**B**) Schematic diagram of niche associated FRC subsets in the LN. (**C**) LN mass and cellularity of CTRL and OXd4. Mann-Whitney test. **P < 0.01; ***P < 0.001. (**D**) Schematic diagram of experimental set-up. (**F**) Immunostaining showing neutrophil distribution (⍺-Ly6G) in CTRL and OXd4 LNs with IgG or ⍺-CD20 treatment. (**E**) Image quantification of neutrophils in the non-draining and draining LN from CTRL and OXd4 mice with IgG or ⍺-CD20 treatment. n = 6-8 mice/group. 2-Way ANOVA test with Šídák’s multiple comparisons test. ***P < 0.001; ns, no significance. (**F**) CFU test quantifying bacterial dissemination to the draining LNs, spleen and blood. n= 4-8 mice/group.

One observation was that the LN was substantially enlarged (**Figure 1C**), with B cells contributing the most to the increase in cell number at OXd4 (**Figure 1C**). When examining the cell distribution within the LN by immunofluorescent (IF) staining and imaging, most of the LN niches appeared enlarged without changing their corresponding niches. However, B cells not only expanded within the B cell zone but also invaded IFZ at OXd4. These changes were associated with a substantial reduction of neutrophil infiltration into the inflamed LN at 4hpi, with the most drastic reduction in the SCS and IFZ in the OXd4 LNs. These changes did not prevent neutrophils from entering the MS (**Figure 1D, E, IgG and Supplemental 1A**).

Based on these observations, we hypothesized that B cells interrupted neutrophil response in the OXd4 LNs. To test this hypothesis, we i.p. injected mice with IgG or ⍺-CD20 and treated mice with OX (**Figure 1D**). We then infected the CTRL or OXd4 mice with *S. aureus* at the lower flank subcutaneously (s.c.) and collected the LNs at 4hpi. Then, neutrophil response in the LN was assessed using IF staining with ⍺-Ly6G antibody. As expected, the neutrophil response was dramatically compromised in the OXd4 LN of mice receiving control IgG (**Figure 1E, F, IgG**). With B cell depletion, neutrophil recruitment was rescued in the OXd4 LN, showing a comparable distribution and number to those in the CTRL LN (**Figure 1E, F, ⍺-CD20**).

To understand whether the restore neutrophil response reduce the possibility of bacterial dissemination, we measured the bacterial load in different organs by counting colony-forming units (CFU) to determine whether rescuing the neutrophil response could prevent bacterial dissemination at OXd4 (**Figure 1D**). With IgG treatment, in CTRL mice, *S. aureus* spread to the draining LNs as early as 4hpi and posed a risk of systemic dissemination to other organs, i.e., spleen, liver, and blood. In OXd4 mice, similar number of the bacteria spread into the draining LNs, with elevated bacterial dissemination into the spleen and blood from 4hpi to 72hpi (**Figure 1G, left**), suggesting a higher risk of bacteremia in mice with OX skin inflammation. In contrast, in OXd4 mice treated with ⍺-CD20 antibody, while *S. aureus* could spread to the draining LN, no bacteria were detected in the spleen or blood (**Figure 1G, right**), indicating that B cell depletion restored neutrophil response and reduced the risk of bacteremia in response to *S. aureus* infection in OXd4 mice.

### OX-inflammation induced a distinct population Cxcl13^int^ RCs

Since immune cell migration and distribution in the LNs are directed by the stromal cells, we hypothesized that remodeling of LN stromal cells resulted in the interrupted neutrophil responses. Flow cytometry analysis showed that FRCs were expanded in the OXd4 LNs compared to the control LN (**Supplemental Figure 1B, C**). To explore the changes in FRC subsets during OX-induced inflammation, we utilized droplet-based scRNAseq on FACS-sorted mouse CD45^−^ cells from the inguinal LNs of CTRL and OXd4 mice (**Supplemental Figure 1D**). After quality control, 3247 cells in the CTRL LNs and 4277 cells in the OXd4 LNs were retained. Next, we performed unsupervised clustering for FRC subsets. UMAP projection identified 11 clusters, each defined by at least 50 marker genes (**Figure 2A**). Consistent with published literature^10^, in the CTRL LNs, we identified Ccl21^+^ TRCs, Cxcl13^+^Cr2^+^ FDCs, Cxcl13^+^Tnfsf11^+^ MRCs, Imnt^+^ SCs, CD34^+^ SCs, and Ccl2^+^Ccl7^+^ MedRCs (**Figure 2B, C**). The Ccl21^+^ TRCs were further split into Slc7a11^+^Ccl19^hi^ (**Ccl19^hi^ TRCs**), Gremlin^+^Ccl19^lo^ (**Ccl19^lo^ TRCs**), and Cxcl9^+^Cxcl10^+^Ccl19^hi^ (**Cxcl9^+^ TRCs**) clusters (**Figure 2B**). While these clusters were conserved in the OXd4 LNs, several significant changes were observed in the FRC subsets on the UMAP (**Figure 2B, C and Supplemental Figure 3**). MedRCs were substantially expanded from 2.19% to 14.96%. There was a significant reduction in Ccl19^lo^ TRCs from 40.61% to 18.17%, accompanied by a substantial spatial transversion of Ccl19^lo^ TRCs on the UMAP, suggesting changes in both the number and their gene expression profile (**Figure 2B, C**). Significantly, a novel Cxcl13^int^Cr2^−^Tnfsf11^−^ cluster (Cxcl13^int^ RCs) was identified in the OXd4 LNs (**Figure 2B, C**), which share no other canonical FRC subset marker genes except the global expression of intermediate level of Cxcl13 compared to FDCs (**Figure 3D, E and Supplemental Figure 4**). As expected, cell proliferation was observed in each FRC subset in the OXd4 LNs, presumably to produce chemokines and create more space to support immune cell expansion in each niche of the LNs (**Supplemental Figure 5, G2m**). However, even after we regressed the cell cycle-related genes^34^ in our dataset, a cluster of dividing cells was still present, characterized by the absence of classic subset signatures (**Figure 2B, C**). Notably, this dividing cell cluster did not show distinct segregation from the CXCL13^int^ RCs on the UMAP and expressed an intermediate level of CXCL13 (**Figure 2D, E and Supplemental Figure 5**), we considered these dividing cells as part of the Cxcl13^int^ RC cluster. Since CXCL13 is known as the most potent B cell chemoattractant^35^, the induction of Cxcl13^int^ FRCs should support B cell expansion in the OXd4 LNs.

**Figure 2.**
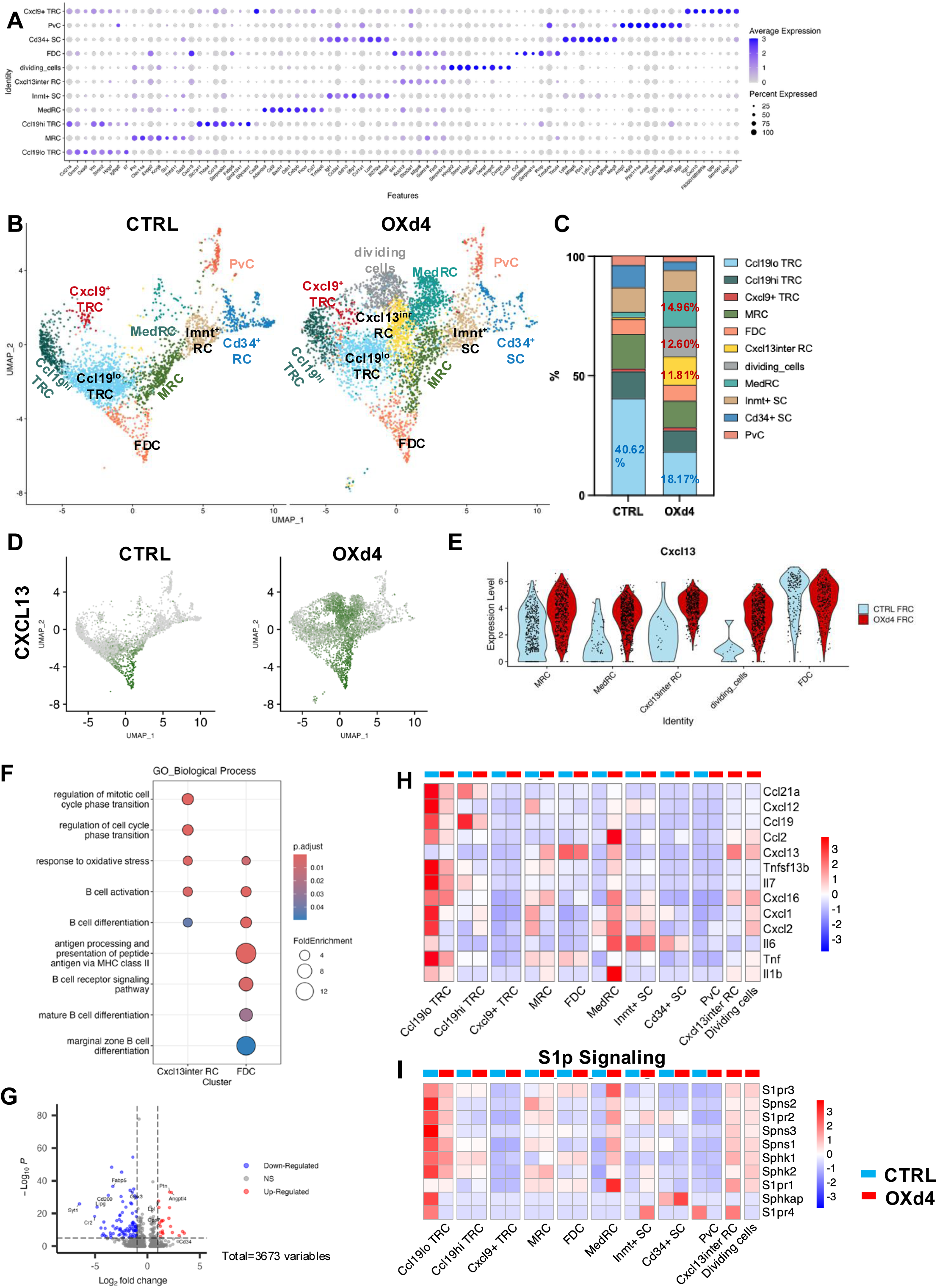
OX-inflammation induced an uncanonical FRC population characterized as Cxcl13^int^ RC. (**A**) Dot plot of the top 8 of most strongly enriched genes for FRC subsets. (**B**) Uniform manifold approximation and projections (**UMAPs**) visualizing FRC clusters from CTRL and OXd4 LNs. (**C**) Stacked bar chart shows the percentage of each cluster in CTRL and OXd4 LNs. (**D**) Expression of Cxcl13 in different FRC subsets. (**E**) Violin charts showing the expression level of Cxcl13 in MRCs, Cxcl13^inter^ RCs, dividing cells, and FDCs. (**F**) Dotplot shows the top 5 biological process signaling pathways that are dominant in Cxcl13int RC or FDC. (**G**) Volcano plot of differentially expressed genes in Cxcl13int RCs compared to FDCs. (**H**) Heatmaps of gene expression of chemokines, cytokines & survival factors. (**J**) Heatmaps of gene expression of S1p signaling.

**Figure 3.**
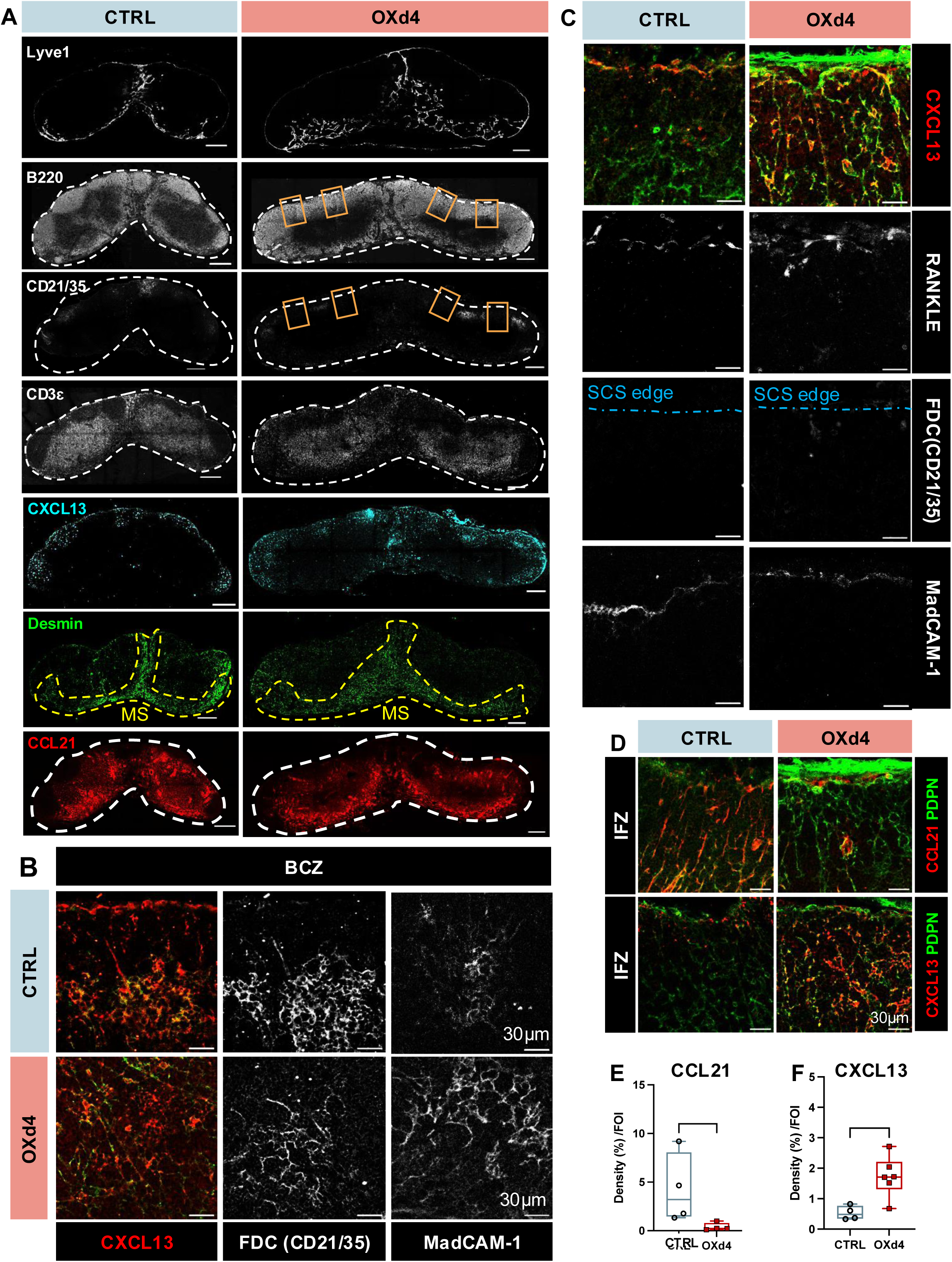
Cxcl13^int^ RCs were mapped to IFZ of the OXd4 LNs. (**A**) Representative tile scan image of the CTRL and OXd4 LNs staining with anti-Lyve1(LECs, MS), anti-B220 (B cells), anti-CD21/35 (FDCs), anti-Desmin (MedRCs), anti-CXCL13 (FDC, MRC and Cxcl13^int^ RCs) and anti-CCL21(TRCs) antibodies. Boxes: IFZs in the OXd4 LNs. n = 5. (**B**) CXCL13 expression in B cell zone (BCZ). FDCs were marked by CD21/35 and MadCAM-1. (**C**) CXCL13 expression in the IFZ. MRCs were marked by RANKL and MadCAM-1 and negative for CD21/35. (**D-F**) Immunostaining and density quantification of CCL21^+^ (**E**) and CXCL13^+^ FRCs (**F**) in the IFZ of CTRL and OXd4 LNs. n=4-6 mice/group. (**E-F**) Mann-Whitney test. *P < 0.05.

FDCs are known as the FRC subsets reside in the B cell zone and the scRNAseq analysis showed they expressed the highest level of Cxcl13 (**Figure 2E**). We used GO analysis to assess the biological pathways in Cxcl13^int^ RCs compared with FDCs and found they were involved in different biological pathways (**Figure 2F**). By differential expression gene (DEG) analysis using the MAST method to compare the transcriptomics between Cxcl13^int^ RCs and FDCs (**Figure 2G**). FDCs are the central component of germinal center (GC) architecture, where they retain immune complexes and present antigens to B cells through their high expression of *Cr2*^36^, *Syt1*^37^, and *Cd200*^38^, which facilitate immune complex trapping, vesicle trafficking, and immune regulation. FDCs are enriched for genes involved in lipid metabolism, including *Lipg* (endothelial lipase)^39^ and *Fabp5* (fatty acid-binding protein 5)^40^, facilitating antigen retention as well as *Gpx3*^41^, supporting long-lived B cell survival. In contrast, Cxcl13^int^ RCs lack these markers, suggesting they do not participate in GC responses. Instead, their expression of *Ptn* and *Angptl4* implies roles in tissue remodeling and stromal support, potentially contributing to the structural integrity of the LN during immune activation. Therefore, Cxcl13^int^ RCs represented a population distinct from FDCs.

Another CXCL13-expressing FRCs are MedRCs in the MS. B cell and plasma cell homeostatic and survival signals (*Cxcl13*, *Tnfsf13b*, *Il6*) were all upregulated in MedRCs of OXd4 LNs (**Figure 2H**), potentially attracted and accommodated more B cells in the MS. Furthermore, MedRCs reside near the exit of LNs where immune cells usually egress from the LN to efferent lymphatic vessels after immune surveillance. Given that lymphocyte emigration from the LN is regulated by S1P-S1PR signaling^42^, we analyzed the S1p signaling-related gene expression, including sphingosine kinases, secretion proteins, and S1P receptors (S1PRs), and found they were upregulated in MedRCs (**Figure 2I**). These data indicated that the expansion of MedRCs increased their ability to attract B cells to the MS and presumably promote cell emigration from the inflamed LNs. However, the biological pathway and the DEG analysis showed Cxcl13^int^ RCs are distinct from MedRCs (**Supplemental Figure 6A, B**). MRCs under the SCS also conservatively express CXCL13. MRCs upregulate Cxcl13 expression in the OXd4 LNs, presumably attract more B cells to the SCS area. Similarly, Cxcl13^int^ RCs showed distinct biological pathways and gene expressional profile compared to MRC, the Cxcl13-expressing FRCs near the SCS (**Supplemental Figure 6A, C**). Thus, Cxcl13^int^ RCs are an induced FRC population distinct from the canonical CXCL13-expressing FRCs.

### CXCL13^int^ RCs are mapped to the IFZs in the OXd4 LNs

To map where the Cxcl13^int^ RCs were located in the LN, we used IF staining to align the FRC subpopulations to the corresponding niches in the LNs. First, by anti-Lyve-1 staining, the lymphatic endothelial cells (LECs) outlined the MS area and showed the expansion of MS in the OXd4 LNs (**Figure 3A, Lyve1**). B cells (anti-B220) were substantially expanded in the BCZ and MS. One notable change in B cell distribution was that B cells had invaded the IFZ of OXd4 LNs (the area between FDC clusters (anti-CD21/35)), replacing the T-cell dominant IFZ, compared to CTRL LNs (**Figure 3A, B220**). T cells (anti-CD3) were expanded in the TCZ and at the border of TCZ and MS, but were diminished in the IFZ (**Figure 3A, CD3**).

To align Cxcl13-expressing FRCs to their corresponding niches, we used anti-CXCL13 staining (**Figure 3A, CXCL13**). As expected, CXCL13 labeled the MRCs and FDCs in the CTRL LN, and CXCL13^+^ signals were dramatically expanded in the OXd4 LN associated with the expansion of B cells. The CXCL13 was highly expressed in the FDCs (CD21/35 labeled) in the B cell zone and the expression was expanded with FDCs and B cells in the OXd4 LNs (**Figure 3A, B, and Supplemental Figure 7A**), suggesting that FDCs maintained their capacity to accommodate B cells within the BCZs. MRCs, labeled by RANKL and MadCAM-1, were located under the SCS above both BCZ and IFZ (**Figure 3A-C**). In OXd4 LNs, CXCL13^+^ FRCs were induced in the IFZ and were negative for FDC marker. Additionally, MRCs remained restricted beneath the SCS (**Figure 3C**). Furthermore, by IF staining with MAdCAM-1, MRCs were restricted in the SCS from day 1-day 4 post OX-treatment (OXd1-OXd4) (**Supplemental Figure 8)**. On the other hand, MedRCs were in the MS area. Desmin is a signature marker of MedRCs, and its expression level did not change between the CTRL and OXd4 LNs (**Supplemental Figure 9A**). By anti-Desmin staining, the expansion of MedRCs was associated with expanded CXCL13 expression in the MS area in the OXd4 LNs (**Figure 3A, Desmin and Lyve-1**). These results suggested that the induced CXCL13-expressing FRCs in the IFZ of OXd4 LNs are the induced Cxcl13^int^ RCs identified by scRNAseq (**Figure 2B**).

IFZ FRCs are Ccl19^lo^ TRCs express CCL21 but not CXCL13 in the control LN (**Figure 3A, D, CCL21**). Staining of CCL21, a cytokine globally expressed by all TRC subsets (**Supplemental Figure 7B)**, showed that in the CTRL LNs, the CCL21^+^ TRCs occupied the TCZs, IFZs, and T-B borders. In the OXd4 LNs, the CCL21^+^ TRCs were still present in the TCZs and T-B borders but sparse in the IFZs (**Figure 3A, D, CCL21**). Correspondingly, while T cells were expanded in the TCZ, the T cell-dominated IFZs in the CTRL LNs were replaced by B cells in the OXd4 LNs (**Figure 3A, B220**). Concomitantly, CCL21^+^Ccl19^lo^ TRCs in the IFZ were diminished with the emerge of CXCL13^+^ FRCs (**Figure 3D-F**). These results suggested that Cxcl13^int^ RCs replaced Ccl19^lo^ TRCs in the IFZ, supporting B cell infiltration into IFZ in the OXd4 LNs.

### Cxcl13^int^ RCs do not compose conduits in the OXd4 LN

In additional to expressing CCL21 to support T cells in the IFZ, Ccl19^lo^ TRCs in the IFZ construct conduits that facilitate lymph communication with HEVs and other cells residing deep in the T cell zone. Since Cxcl13^int^ RCs replaced the Ccl19^lo^ TRCs in the IFZ, we assessed the biological pathways in Cxcl13^int^ RCs compared to Ccl19^lo^ and Ccl19^hi^ TRCs (**Figure 4A**). Cell chemotaxis was detected in both Cxcl13^int^ RCs and TRCs clusters. B cell activation and cellular response to fibroblast growth factor stimulus were found in Cxcl13^inter^ RCs. However, collagen fibril and extracellular matrix organization, were not detected in the Cxcl13^int^ RCs (**Figure 4A**), suggested that Cxcl13^int^ RCs were poor at constructing conduits. Furthermore, we assessed the gene expression of conduit components and ECM remodeling proteins (**Supplemental Figure 9C**). In CTRL LNs, the significant populations producing conduit components and ECM remodeling proteins were Ccl19^lo^ TRCs, Ccl19^hi^ TRCs, MRCs, and Cd34^+^ SCs. In the OXd4 LN, the gene level of ECM components and remodeling protein was comparable except significantly elevated in the MedRCs. However, Cxcl13^int^ RCs produced lower conduit component genes than Ccl19^lo^ TRCs but with a relatively high level of ECM modifying protein, i.e., *Mmp9* and *Mmp23* (**Supplemental Figure 9C**).

**Figure 4.**
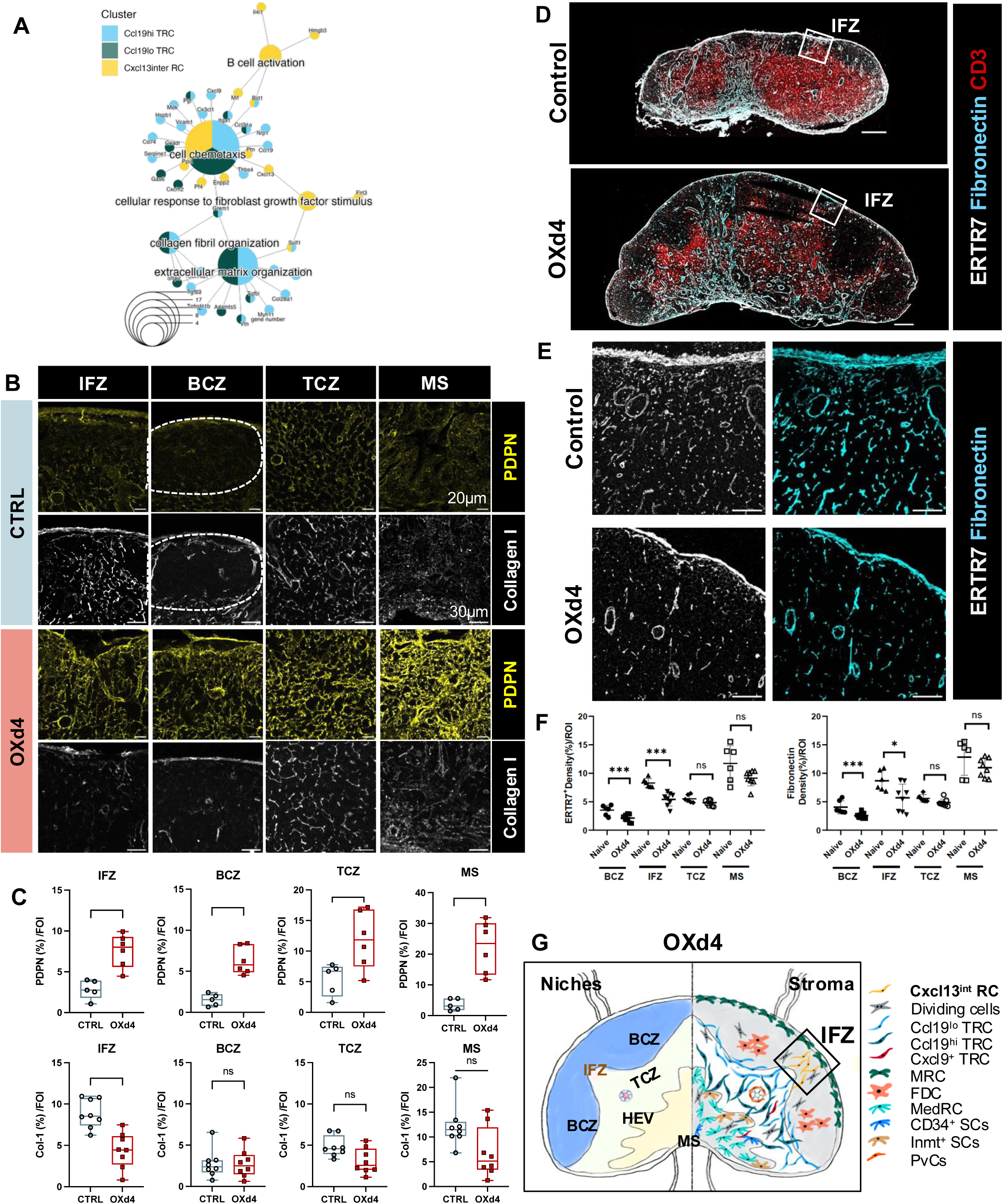
Cxcl13^int^ RCs compromised the conduit network in the IFZs of the OXd4 LNs. (**A**) CnetPlot showing gene ontology (**GO**) analysis of the representative biological process in Ccl19^hi^ TRC, Ccl19^lo^ TRC, and Cxcl13^int^ RC clusters. (**B**) Immunostaining of conduit (grey) and FRC (yellow) in different LN regions, n=6 mice/group. (C) Image quantification of PDPN+ and Collagen I density per field of interest (**FOI**) in different niches of LN comparing CTRL and OXd4 (Bottom), n=5-8 mice/group. Mann-Whitney test. *P < 0.05; **P < 0.01; ns, no significance. (**D**) Tile scans of CTRL and OXd4 LNs with anti-ERTR7 (grey), anti-Fibronectin (cyan), and anti-CD3 (red) staining. (**E**) Zoom-in images of IFZ in the CTRL and OXd4 LNs with anti-ERTR7 (grey) or anti-Fibronectin (cyan) staining. (**F**) Quantification of images from (B). 2-Way ANOVA test with Šídák’s multiple comparisons test. *P < 0.05; ***P < 0.001; ns, no significance. (**G**) Schematic Figure of niches associated FRCs in the OXd4 LN.

To validate the scRNAseq analysis, we performed IF staining to map the changes in FRC-conduit network using anti-PDPN and anti-Collagen I. PDPN^+^ FRCs were expanded in all the niches in the OXd4 LNs. B cell zones are known to have sparse conduits. Only IFZ showed significant reduction in Collegen I^+^ conduits (**Figure 4B**). The conduit reduction in the IFZ was consistent when using ER-TR7 and fibronectin (**Figure 4C, D and Supplemental Figure 10**). Thus, when the poor conduit-producing Cxcl13^int^ RCs replaced the potent conduit-forming Ccl19^lo^ TRCs in the IFZ, FRC remodeling led to increased CXCL13 to recruit B cells, instead of expressing CCL21 to attract T cells or forming conduits as seen in the control LNs (**Figure 4E**).

### The induction of Cxcl13^int^ RCs in the OXd4 LN depends on B cells

Generally, the FRCs produce chemokines to coordinate the spatial arrangement of immune cells. In turn, immune cells also modulate the FRC differentiation and phenotype^14,43–45^. Therefore, we hypothesized that Cxcl13^int^ RCs were induced by interacting with immune cells. We ran an interactome analysis using CellChatDB by combining a dataset of CD45^+^ immune cells isolated from immunized LN^14^ with our FRC dataset. The immune cells and each subset of FRCs exhibited complicated interactions in both the control and OXd4 conditions, but only B cells were found to interact with Cxcl13^int^ RCs at OXd4 (**Figure 5A and Supplemental Figure 12**). Therefore, we hypothesized that B cells induced Cxcl13^int^ RCs in the inflamed LN.

**Figure 5.**
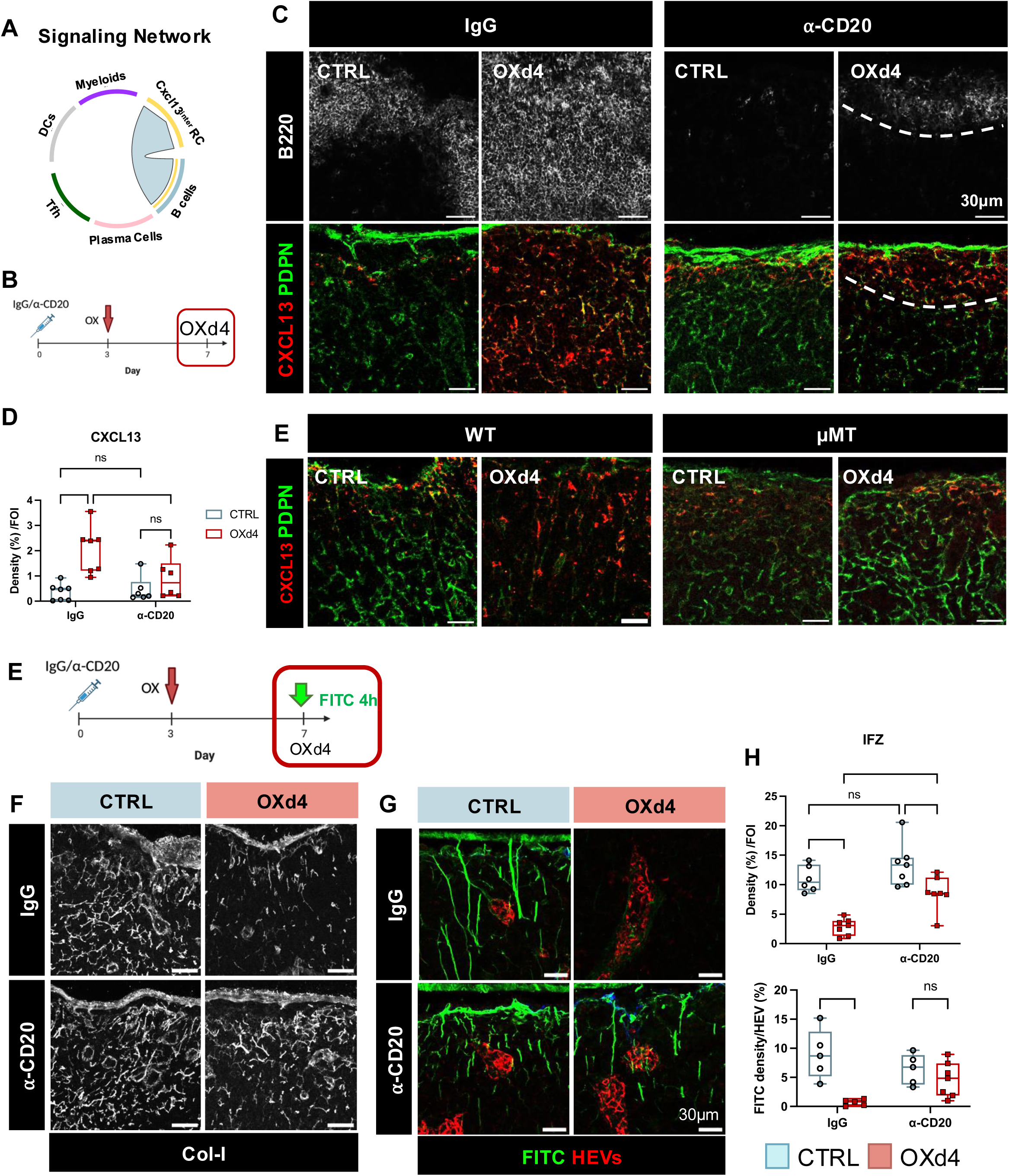
B cell depletion eliminates Cxcl13^int^ RCs and retains conduit-mediated lymph drainage to the HEVs in the inflamed LN. (**A**) Interactome analysis in signaling pathways using CellChatDB showed only B cells interact with Cxcl13^int^ RCs (Immune cell dataset is from M Lütge *et al*., 2023). (**B**) Schematic diagram of experimental design. (**C**) Immunostaining showing B cells (⍺-B220) and Cxcl13^+^ RCs (⍺-Cxcl13,⍺-PDPN) in CTRL and OXd4 LNs with IgG or ⍺-CD20 treatment. (**D**) Quantification of Cxcl13^+^ RC density in (**C**). n = 6-7 mice/group. 2-Way ANOVA test with Šídák’s multiple comparisons test. *P < 0.05; ***P < 0.001; ns, no significance. (**E**) Schematic diagram of experimental design. (**F-H**) Conduits and conduit-mediated lymph drainage to HEVs. (**F, G**) Representative image and (**H**) quantification of conduit network (⍺-Collagen I) and conduit-mediated FITC drainage to HEVs in CTRL and OXd4 LNs with IgG or ⍺-CD20 treatment. (**F**) n=6-7 mice/group and (**G**) n=5 mice/group, 2-Way ANOVA test with Šídák’s multiple comparisons test. *P < 0.05; **P < 0.01; ***P < 0.001; ns, no significance.

To test this hypothesis, we utilized an anti-CD20 antibody to deplete B cells in WT mice via intraperitoneal (i.p.) injection. We applied OX to induce skin inflammation three days post-injection (to enable maximum B cell depletion) (**Figure 5B**). The depletion can persist for 21 days^46^ but retained about 0.3% of B cells in the LN (**Supplemental 13A-B**). Our results showed that the depletion of B cells successfully eliminated Cxcl13^int^ RCs in the IFZ (**Figure 5C-D**). Since anti-CD20 could not completely abrogate B cells, a few CXCL13^+^ RCs were highly correlated with the residual B cells (**Figure 5C, ⍺-CD20 OXd4**). To rule out the impact of the residual B cells, we additionally used μMT mice, a B cell-deficient mouse model, and applied OX to induce skin inflammation. No expansion of Cxcl13^int^ RCs was detected in the OXd4 LN of μMT mice (**Figure 5E**). Although CXCL13^+^ RCs were detected in the LN of μMT, they should be classified as MRCs since they are MadCAM-1^+^ (**Supplemental Figure 12C**). These results indicate that MRCs are independent of B cells in both control and OXd4 LNs, whereas B cells are required to induce Cxcl13^int^ RCs in the inflamed LN.

### B cell depletion preserves conduit network and lymph drainage in the LN during OX-induced inflammation

Having demonstrated that depleting B cells eliminated Cxcl13^int^ RCs in the OXd4 LN, we examined if we could restore the conduit integrity and lymph drainage along the network to HEVs. We used ⍺-CD20 antibody to deplete B cells and then applied OX (**Figure 5E**). Using ⍺-collagen I antibody, we imaged and quantified the density of the conduit network in the IFZs. Compared to the IgG control group, B cell depletion did not significantly change the conduit network in CTRL LNs but preserved the integrity in the OXd4 LN (**Figure 5F, H**).

Next, we applied FITC as a lymph tracer and assessed the lymph drainage in the LN and the amount reaching the HEVs. B cell depletion restored the lymph drainage in the OXd4 LN and when focusing on the IFZs, the drainage to HEVs was also rescued in the OXd4 LN with B cell depletion (**Figure 5G, H and Supplemental Figure 13**). Collectively, these results showed that B cell depletion retained the integrity and function of the conduit network in the OXd4 LNs.

### B cell-derived lymphotoxin signaling induces Cxcl13^int^ RCs in the inflamed LN

Finally, we aimed to determine the molecular mechanism by which B cells induce Cxcl13^int^ RCs. Using interactome analysis, we plot the ligand-receptor communication between Cxcl13^inter^ RCs and B cells. When Cxcl13^int^ RCs are a source of ligands, there is a multitude of interactions towards B cells (source of receptors), including chemotactic signals, B cell differentiation, and antigen presentation (**Figure 6A**). However, the only interactions from B cells (ligands) towards Cxcl13^int^ RCs (receptors) are lymphotoxin pathways (**Figure 6B**). LTβR and TNF receptor 1 are widely expressed by all FRC subsets and comparable between CTRL and OXd4 LN (**Supplemental Figure 14**). Lymphotoxins (membrane-bound LT⍺_1_β_2_ and secretive LT⍺_3_) are expressed by B cells in homeostasis and can be upregulated by interacting with CXCL13^47^ or during GC formation^48^. To determine whether B cells induced Cxcl13^int^ RCs via lymphotoxin signaling, we adoptively transferred B cells isolated from WT and LT⍺^−/-^mice to the μMT mice. Four weeks later, giving sufficient time for B cell reconstitution in the LN, mice were treated with OX to induce inflammation (**Figure 6C**). By IF staining, in the CTRL LNs, CXCL13^+^ MRCs were detected in the SCS in both WT and μMT mice (**Figure 6D**). Cxcl13^int^ RCs expanded in the IFZ of WT but not μMT in OXd4 (**Figure 6D, E**). Without the OX treatment, Cxcl13^+^ MRCs were found in the SCS of μMT that received either WT or LT⍺^−/-^B cells. In the OXd4 LNs, the μMT mice that received WT B cells had an expansion of Cxcl13^int^ RCs. However, the μMT mice receiving LT⍺^−/-^B cells did not induce Cxcl13^int^ RCs in the OXd4 LNs (**Figure 6D, E**). Taken together, these data showed that Cxcl13^int^ RCs were induced by B cell-mediated lymphotoxin signaling in the inflamed LN (**Figure 6F**).

**Figure 6.**
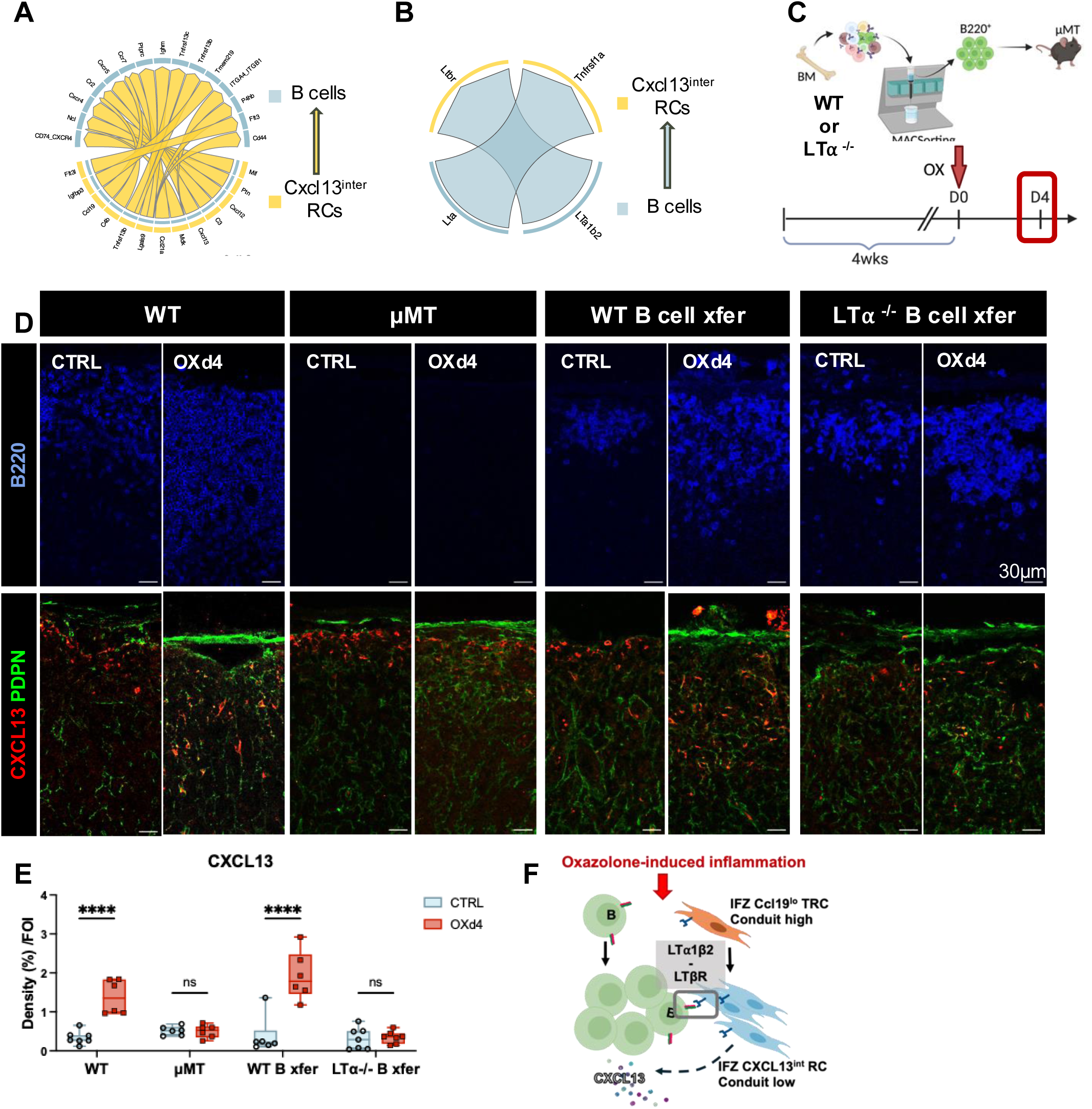
B cell-derived lymphotoxin signaling induces Cxcl13^int^ RCs in the inflamed LN. (**A, B**) Specific pathways of B cells and Cxcl13^int^ RCs interactome analysis. (**C**) Schematic diagram of B cell transfer experimental design. (**D**) Representative images and (**E**) quantification of Cxcl13^+^ RCs (anti-Cxcl13, anti-PDPN) by IF in WT, muMT, WT B cell transfer, and LT⍺^−/-^B cell transfer groups. n=6-7 mice/group. 2-Way ANOVA test with Šídák’s multiple comparisons test. ****P < 0.0005; ns, no significance. (**F**) Schematic diagram showing the induction of Cxcl13^int^ RCs by B cells via lymphotoxin signaling in the LN during OX-induced inflammation.

### Cxcl13^int^ RCs display a transitional phenotype during OX-inflammation

Furthermore, the scRNAseq velocity analysis showed that Cxcl13^int^ RCs display a vector towards the CCL21^+^Ccl19^lo^ FRCs, suggesting this inducible population might differentiated into IFZ FRCs later after the OX-inflammation (**Figure 7A**). To test this possibility, we collected LNs from mice at 7- and 14-days post-OX treatment (OXd7 and OXd14) and found IFZ FRCs regained the CCL21 and reduce CXCL13 expression over time in the IFZ (**Figure 7B)**. Concomitant with the revert of Cxcl13^+^ FRCs to CCL21^+^ FRCs in the IFZ, the conduits (Col I) and conduit-mediated lymph drainage to HEVs were also restored over time from OXd7-OXd14 (**Figure 7C**). Finally, neutrophil response to *S. aureus* at 4hpi were also restored over time from OXd7-OXd14 (**Figure 7D)**. These results suggest that B cell-induced Cxcl13^int^ RCs represent a transitional phenotype, creating a time window of compromised neutrophil responses.

**Figure 7.**
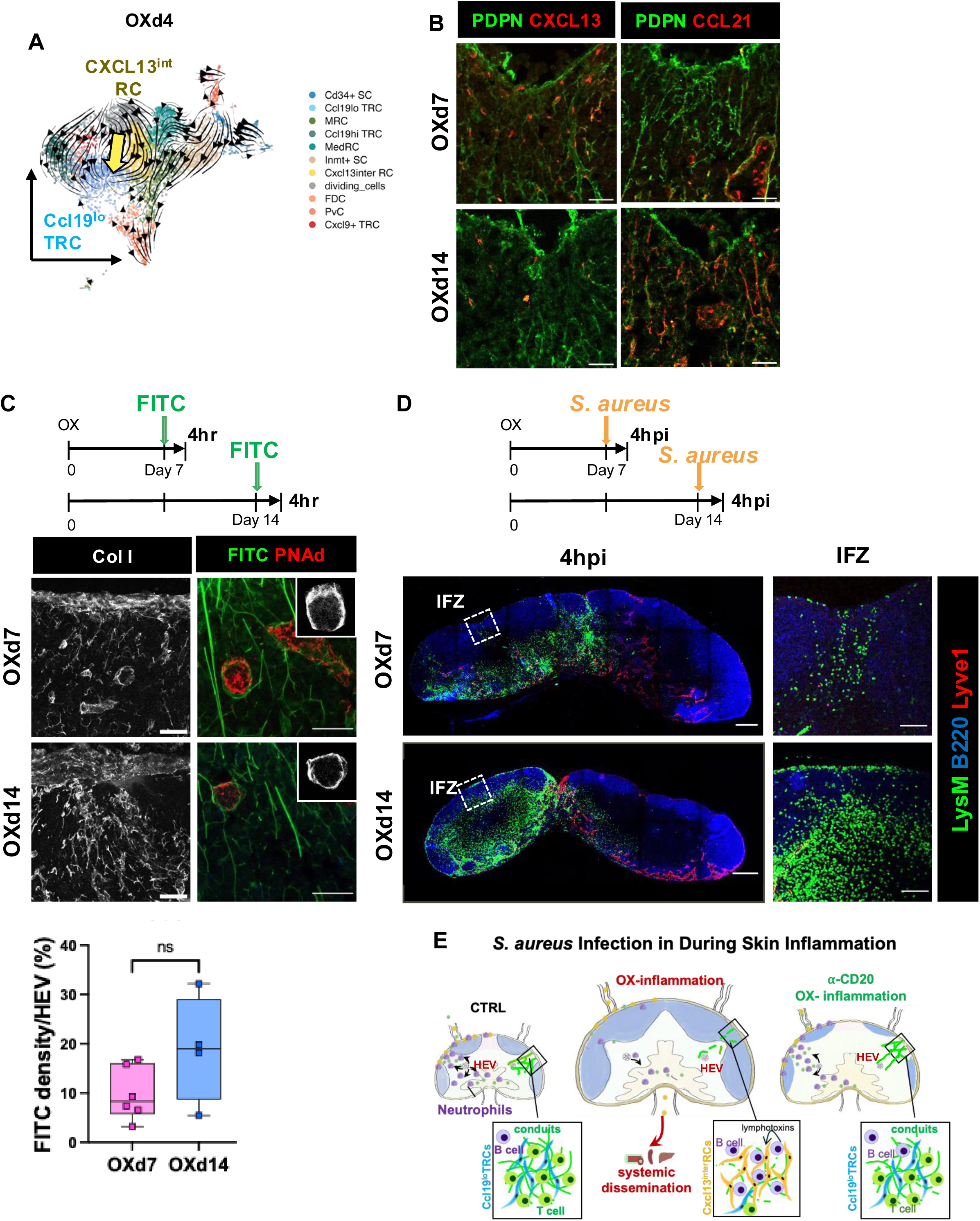
The induced Cxcl13^int^ RCs display transitional phenotype during OX-inflammation. (**A**) Velocity analysis suggested Cxcl13^int^ RC could be converted to Ccl19^lo^ TRCs. (**B**) Immunofluorescence images of CXCL13^+^ FRCs, CCL21^+^ FRCs in the IFZ at OXd7 and OXd14. (**C**) Col-1^+^ conduits and FITC drainage to HEVs in the OXd7 and OXd14 LNs. Image quantification of FITC drainage into HEVs. n=4-6 mice/group Mann-Whitney test. *P < 0.05; ns, no significance. (**D**) Immunofluorescence images showing neutrophil distribution (LysM^+^, green) at 4 hours post-S. aureus infection in OXd7 and OXd14 LNs. (**E**). Schematic summary. In healthy conditions, *S. aureus* infection rapidly recruits neutrophils to the LN. OX-inflammation remodels FRC-conduit network and impairs the neutrophil response to *S. aureus* infection, posing a higher risk of bacteria systemic dissemination. B cell depletion preserves FRC-conduits and rescues the neutrophil response to infection during OX-inflammation.

## Discussion

This study investigated how FRC subsets were remodeled during OX-induced inflammation and how the remodeled FRC network interrupted neutrophil response to the secondary *S. aureus* infection during OX-inflammation. scRNAseq analysis showed that nearly every population of FRCs showed changes in cytokines/chemokine expression profile and some level of proliferation without changing their signature genes. Notably, a new FRC subset, Cxcl13^int^ RCs emerged in the IFZ of the inflamed LN. This FRC subset was induced by B cells, replaced the canonical IFZ TRCs, disrupted the IFZ conduit network, and compromised conduit-mediated lymph drainage to HEVs (**Figure 7E**). Our study highlights the crucial role of the B cells in regulating FRC network remodeling during OX-inflammation.

It has been recognized that FRC-conduit network mediated lymph-borne small molecular weight factors (free-form antigens, cytokines and chemokines) draining into the deep LN paracortex, and the IFZ FRC-conduit network plays a crucial role in guiding lymph-HEV communication^19,21,23^. LNs play important roles in preventing bacteria systemic dissemination and the innate immune responses were essential for the early protections^5,6,24^. Furthermore, the importance of conduit-mediated lymph drainage in regulating neutrophil infiltration in the LNs in response to *S. aureus* infection. In the OX-induced inflammation, when Cxcl13^int^ RC replaces Ccl19^lo^ TRCs in the IFZ, conduit network integrity and lymph drainage to HEV were interrupted. Eliminating Cxcl13^int^ RCs via B-cell depletion preserved the conduits, retained conduit-mediated lymph drainage and rescued neutrophil response to *S. aureus* infection during skin inflammation (**Figure 7E)**.

Several studies have revealed FRC heterogeneity in lymphoid organs in mice and humans using scRNAseq^10–13,31,32^. The FRC subsets discovered in CTRL LN from our dataset are consistent with studies by LB. Rodda *et al*.^10^, VN. Kapoor *et al*.^11^, and J D’Rozario *et al.*^32^. However, the Ludewig group^13,31^ detected a few additional sub-clusters of TRCs, i.e., TBRCs (*Ccl21a, Gremlin, Fmod*) and IFRCs (*Hamp2, Cavin2*). These marker genes were detectable in Ccl19^lo^ TRC and MRC from our dataset, but we did not have the cellular resolution to accurately discern distinct sub-clusters. The gene expression profiles of FRC subsets under inflammatory conditions have been assessed using scRNAseq at 3 days post-LCMV infection^31^, day 12-14 post-VSV infection^13,14^, and day 3 post-LPS stimulation^32^. The studies using virus infection models focused on examining B cell-associated FRCs and the crucial role of FRCs in regulating B cell recruitment, proliferation, and germinal center formation, while the LPS stimulation showed the importance of FRCs in regulating LN resident macrophages. Although global alterations in the gene profile were detected, no new clusters were reported in those stimulations. Thus, the induction of Cxcl13^int^ RCs may be due to different types of immune stimulation.

Observation of non-classical CXCL13^+^ SC population has been reported in the LN 3 weeks after CFA-induced inflammation^49^. In this model, B cells trespassed into TCZs, converting TRCs at the border of T and B cell zones into CXCL13-expressing cells, setting up new boundaries of B cell follicles^49^. In another study using Helminth infection in the gut, the induction of CXCL13^+^ MRC-like cells populating the IFZs were observed to supporting B cell and GC formation in the mesenteric LNs (mLNs) at 21 days post-infection and reported ^50^. It is unknown whether the Cxcl13^int^ RCs identified in this study are the same or distinct from those CXCL13^+^ SCs. However, the Cxcl13^int^ RCs identified in this study only show their supporting B cell expansion to IFZ, but not GC formation (**Figure 2F, G**). Furthermore, Cxcl13^int^ RCs displayed a transitional phenotype at OXd4, likely to support B cell expansion at the initial phases of immune response and could be revert to canonical Ccl19^lo^TRCs in the IFZ when the OX-inflammation was resolved. These findings suggest that LN stromal cell remodeling is dynamically regulated over the course of an immune response. Thus, non-canonical Cxcl13^+^ RCs may be induced under different conditions but are only detectable within specific time windows.

Additionally, the induction of CXCL13-expressing FRCs in CFA and Helminth models both required lymphotoxin-dependent interaction, as evidenced by transferring LTβ^−/-^B cells to μMT mice^49,50^. These studies are consistent with our study transferring LTα^−/-^B cells to μMT mice, demonstrating the induction of CXCL13^int^ RCs relies on B cell-mediated lymphotoxin signaling. Moreover, *in vitro* experiments proved that engagement of LTβR promotes CXCL13 expression in the LN stromal cell line^51^ and lung fibroblast line^52^. A depletion of LTβR in FRCs impedes B cell follicle formation in mLNs following intestinal Helminth infection^29^. These studies are also consistent with our observation in the OX-inflammation, where B cells induce CXCL13^int^ RCs and, in turn, attract more B cells infiltrating via the production of CXCL13 (**Figure 5**). Since LTβR signaling is essential for maintaining and regulating LN architecture, although we cannot exclude the possibility that B cell-derived LTα_3_ may play a role in inducing CXCL13^int^ RCs, it is more likely that this process is through LT⍺_1_β_2_ -LTβR signaling.

FRCs play a crucial role in building the conduit network. The disrupted conduit network in the inflamed LN have been reported in various inflammatory models^7,22,24,33^. The reduced conduits were most drastic in the IFZ than other regions (**Figure 4 and Supplemental Figure 10**) The molecular mechanism of conduit alteration has only been studied using FRC culture before. Martinez *et al.* showed that ECM secretion was inhibited due to reduced expression of transporter protein LL5β after PDPN-CLEC-2 interaction^7^. Studies on other cell lines indicate a possible mechanism of why it reduced the ability of conduit maintenance. Treating with LTα in human nucleus pulposus cells^53^ and chondrocytes^54^ increased matrix degrading, including MMP3, MMP9, and MMP13, and decreased ECMs, such as type II collagen and aggrecan. Our study showed that the poor-conduit producing Cxcl13^int^ RCs replaced the potent conduit-making Ccl19^lo^ TRCs in the IFZ, resulted in the reduction in conduit network in IFZs of OXd4 LNs. Our studies suggest that B cell-mediated LT signaling is essential for the induction of Cxcl13^int^ RCs and the disruption of conduits as shown by the scRNAseq analysis and validated by transferring WT or LTα^−/-^B cells to B cell-deficient μMT mice.

In summary, our study provides a mechanism by which FRC-conduit network is remodeled in OX-skin inflammation. Since OX-induced inflammation is often used to model atopic dermatitis or contact hypersensitivity, our studies provide the potential mechanism to interpret why dermatitis patients are prone to *S. aureus* infection and have a higher risk of bacteremia. Our study also provides evidence that B cell-stromal interaction might be a potential target for improving host immunity to secondary infections.

### Limitation of the Study

The Cxcl13^int^ RCs replaced of Ccl19^lo^ FRCs in the IFZ, indicating the Ccl19^lo^ TRC cluster shift observed on the UMAP of scRNAseq analysis might result from to changes in the niches of the LNs. However, the origin of Cxcl13^int^ RCs remains unclear. On the other hand, this study provided evidence that LN stromal remodeling interrupted the neutrophil response to *S. aureus* during OX-inflammation. However, the initial wave of neutrophil infiltration represents an early step in the host immune defense. Other immune cells would be activated to combat the infection. How LN stromal cell remodeling impacts other immune cells in the LN during OX-inflammation and in other diseased conditions remains to be investigated. Furthermore, how the LN stromal cell remodeling during OX-inflammation in mice aligns with skin diseases or other human pathologies also requires further studies.

## Methods and Materials

### Mice

C57BL/6 mice were purchased from the Jackson Laboratory. μMT mice breeders were purchased from the Jackson Laboratory and bred at the Health Sciences Animal Resource Center at the University of Calgary. 6-12 weeks old male and female mice were used for all experiments, except scRNAseq, in which only female samples were used. All animal protocols were reviewed and approved by the University of Calgary Animal Care and Ethics Committee and conformed to the Canadian Council on Animal Care guidelines.

### Oxazolone Skin Sensitization

To induce skin and LN inflammation, 200μL of 4% Oxazolone (OX, Sigma) suspended in acetone was applied topically on the shaved abdomen. Inguinal LNs (the draining LNs) were collected at time points of interest.

### Single-cell RNA Sequencing (scRNAseq) Sample Preparation

Inguinal LNs were harvested from naïve mice (n=5) and OXd4 mice (n=4) and pooled to get enough SCs. LNs were pierced into small pieces (∼1 mm^2^) using fine forceps and placed in 5ml of RPMI-1640 on ice. Once all LNs were processed, RPMI-1640 was replaced by enzyme cocktail of RPMI-1640 containing 2% FBS, 0.8 mg/ml Dispase II, 0.2mg/ml Collagenase P, and 0.1 mg/ml DNase I. Tubes were incubated at 37°C in a water bath for 15min in total. Tubes were gently inverted every 5 mins to ensure efficient digestion. Then, the mixture was pipette up and down until obvious fragments were gone. Cells were then filtered through 40μm -filter cap tube, washed with 2% BSA and counted. LN SCs were enriched using CD45 microbeads and autoMACS system (Miltenyi). We used 7μl beads per 10^7^ cells, incubating in 2% BSA buffer for 15min in dark at 4°C. Cells were washed once and unlabelled portion was collected following the manufacturer’s instructions. Enriched SCs were then counted and stained for L/D, CD45, Ter119 for FACS sorting. Live CD45^−^Ter119^−^LN SCs were then sorted to high purity using FASAria with 100 um tip at 20psi.

### scRNAseq library construction, sequencing, and analysis

Single cells from each sample were loaded for partitioning using 10X Genomics NextGEM Gel Bead emulsions (v3.1). Each sample was processed according to the manufacturer’s recommended protocol (PCR amplification steps were run at 12X and 14X, respectively). Final cDNA library size determination and QC were performed using TapeStation D1000 assay. Sequencing was performed using Illumina NovaSeq S2 and SP 100 cycle dual lane flow cells over multiple rounds at the UCalgary Centre for Health Genomics and Informatics (CHGI). 11,126 and 13,348 cells were captured and sequenced to approximately 30, 824 and 24,060 reads per cell in the CTRL and OXd4 LNs, respectively. Sequencing reads were aligned using CellRanger 6.1.2 pipeline^55^ to the standard pre-built mouse reference genome (mm10-3.0.0).

Downstream analyses were performed using the Seurat package^56^ running in R. FRC were pre-filltered using Ptprc (=0), Cdh5 (=0), Prox1 (=0), Lyve1 (=0), and Pdgfra (>1). PvC were pre-filtered using Itga7(>-1). FRC and PvC were then merged and performed quality control including excluding cells with less than 500 or more than 5000 measured genes, >10% mitochondrial counts, and <7% ribosomal genes. The data were then normalized and performed FindVariableFeatures function using “vst” method in the space of 2,000 most variable genes. Subsequently, the CTRL and OXd4 datasets were integrated, scaled with regression of cell cycle score, percentage of mitochondrial genes and ribosomal genes, dimensional reduction with principal component analysis (PCA). 1-30 principle components were input for FindNeighbors based on the JackStraw method. Clusters of cells were identified with graph-based clustering (FindNeighbors, resolution = 0.8). Dimensional reduction via UMAP was then performed with providing 1-30 principal components. Clusters of cells were denoted by running FindConservedMarkers function (min.pct = 0.1, logfc.threshold = 0.25, test = “MAST”).

Heatmaps were generated using AggregatedExpression function in Seurat and pHeatmap package. GO analysis were performed using ClusterProfiler package. Cell interactome analysis with *M Lütge et al.,* dataset was performed using CellChatDB package.

RNA velocity analysis were performed in Python with scVelo package. To generate the loom file for each sample, we compiled and ran velocyto using the filtered_feature_bc_matrix/barcodes.tsv files, the possorted_genome_bam.bam files of each sample and the reference genome. This outputted a .loom file for each sample. Then, all loom files from all samples were combined to generate one final loom file to proceed with downstream analysis with scVelo.

### B cell depletion using anti-CD20 antibody

200μg Rat IgG or ⍺-CD20 was intraperitoneally (i.p.) injected into mice. Three days later, mice were treated with OX topically on the shaved abdomen.

### B cell transfer

Firstly, B220+ cells were isolated from bone marrow (BM) of C57/BL6 or LT⍺^−/-^ mice using B220 beads and MACS sorting (Miltenyi). 10^7^ cells were adoptively transferred to μMT mice via tail vein 4 weeks later, mice then were treated with OX on shaved abdomen and samples were collected 4 days later for downstream experiments.

### Tracing Lymph Drainage by FITC

100μL 2% FITC (Sigma) in 1:1 (v/v) acetone/dibutyl phthalate mixture was applied topically on the shaved lower flank. Inguinal LNs were collected 4 hours post-treatment.

### Methicillin-resistant *Staphylococcus aureus* (MRSA) Infection

A community-acquired MRSA, strain MW2 (USA 400), was a kind gift from Dr. Paul Kubes’ lab. The bacteria were cultured overnight at 37°C by tipping the frozen stock to a 15mL Falcon tube containing 5mL Brain-Heart-Infusion (BHI) culture media to make a primary culture. Then, 500μL of primary culture was added to a conical flask containing 50mL BHI and cultured at 37 degrees in a shaking incubator for 2h30min. The Optical Density (OD_660nm_) was measured, which should be around 0.7 (e8 CFU/mL). 2.5×10^7^ *S. aureus* resuspended in a 50μL sterile PBS were injected intradermally (i.d.) in the lower flank of the mice.

### Colony-Forming-Unit (CFU) Test

4, 24, and 72 hours post-infection, samples of interest were collected using a sterile technique. Blood was collected into a heparinized Eppendorf by retro-orbital. 50μL blood was plated onto BHI agar plates. The inguinal LNs (draining LNs), axillary LNs (downstream LNs), spleens, livers, abdomen skins, and infected flanks were harvested, weighed, and homogenized. LNs were homogenized using a VWR Pellet Mixer in 300μL of sterile PBS. Other tissues were homogenized using Kinematica PT10-35 Homogenizer Mixer. 100μL tissue homogenate with serial dilutions was plated onto BHI agar and incubated at 37°C for 15h, and then bacterial colonies were counted.

### Immunofluorescent Staining

LNs were harvested, fixed with 4% paraformaldehyde (PFA) overnight, and transferred to 30% sucrose until sinking to the bottom. Next, LNs were embedded in OCT and frozen on a dry ice/ethanol mixture. For some specific stainings (i.e., CD3e, Collagen I, ERTR7), LNs were embedded in OCT right after the collection. OCT blocks were then sectioned in 10 μm using a cryostat (Leica).

Sections were washed once with PBS to remove OCT and blocked with 5% mouse serum for 1h. Next, primary antibodies of interest were added and incubated overnight at 4 degrees. After three washes with PBS, sections were incubated with secondary antibodies for 1h at room temperature. The sections were washed three times with PBS and mounted with mounting media with DAPI (Product#). For staining, including biotin, an extra biotin/strep blocking (Biolegend#) was performed after the 5% mouse serum blocking.

Antibodies and dilutions used in this study are provided in Supplemental Table 1.

### Flow Cytometry

LNs were gently pressed through a 40-μm strainer to create a single-cell suspension and then washed with FACS buffer (1% bovine serum azide (BSA), 0.1% sodium azide in PBS). The cells were blocked using anti-CD16/32 antibody for 5min at room temperature, then stained with Fixable Viability Dye for 30min at 4 degrees. The cells were washed with FACS buffer and stained with conjugated antibodies of interest for 30min at 4 degrees. After washing with FACS buffer, the cells were fixed with 2% PFA in FACS buffer. If using unconjugated antibodies, primary and secondary staining were performed after live/dead staining. The cells were acquired within a week by BD FACSCanto, and data were analyzed using FlowJo. The antibody list is provided in Supplemental Table 2.

### Microscopy

Fluorescent images were taken with a SP8 confocal microscope (Leica). The objectives were 10x, 25x in air, and 63x in oil for different purposes. Image processing, adjustment, and quantification were performed using the ImageJ software.

### Statistics

The data were expressed as mean *±* standard deviation (SD). Quantitative data were tested using indicated statistical tests in the GraphPad Prism software. *P < 0.05; **P < 0.01; ***P < 0.001; ****P < 0.0005; ns, no significance.

## Acknowledgement

This study is supported by Canadian Institutes of Health Research project grant, and the Dianne and Irving Kipnes Foundation (SL). In memory of Dr. Dianne Kipnes, whose philanthropic support for lymphedema and lymphatic research through the Dianne and Irving Kipnes Foundation and invaluable contributions as a lymphedema patient involved in lymphatic studies will always be remembered.

