## Supplemental Figures for "B Cell-Induced Lymph Node Stromal Remodeling Compromised Neutrophil Response to Secondary *Staphylococcus aureus* Infection"

**Supplemental Figure 1**

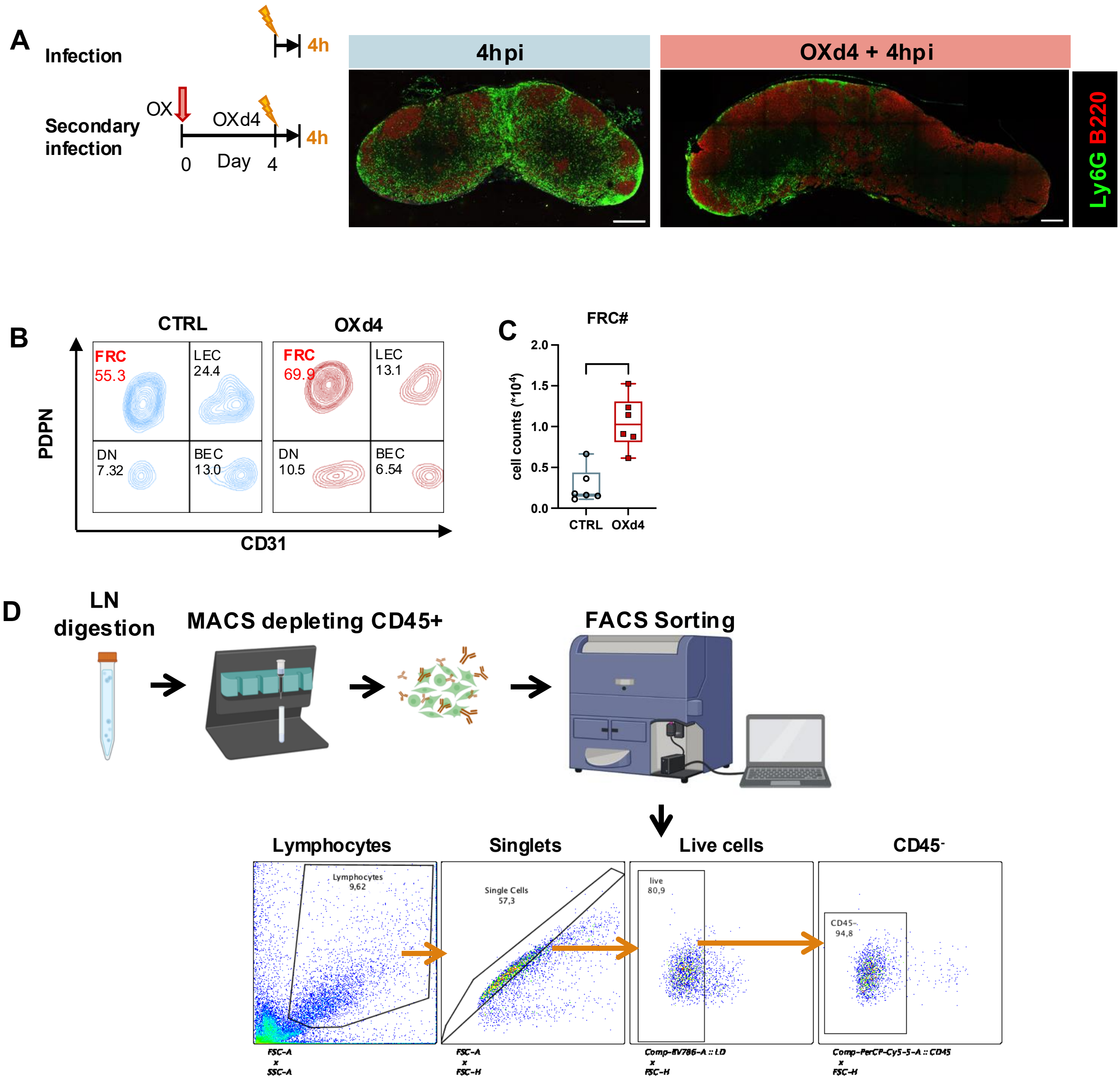


(**A**) Immunostaining showing neutrophil distribution (α-Ly6G) in CTRL and OXd4 LNs at 4hour post-infection (4hpi) to S. aureus. **B**) Representative contour plots of stromal cell gating in the CTRL and OXd4 LNs. (**C**) Flow cytometry analysis of LEC number ($CD45^{-}PDPN^{+}CD31^{+}$) and FRC numbers ($CD45^{-}PDPN^{+}CD31^{-}$) and in CTRL and OXd4 LNs. Mann-Whitney test. *P < 0.05; **P < 0.01; ns, no significance. (**D**) Schematic diagram of LN digestion and sorting strategy for scRNAseq preparation.

**Supplemental Figure 2**

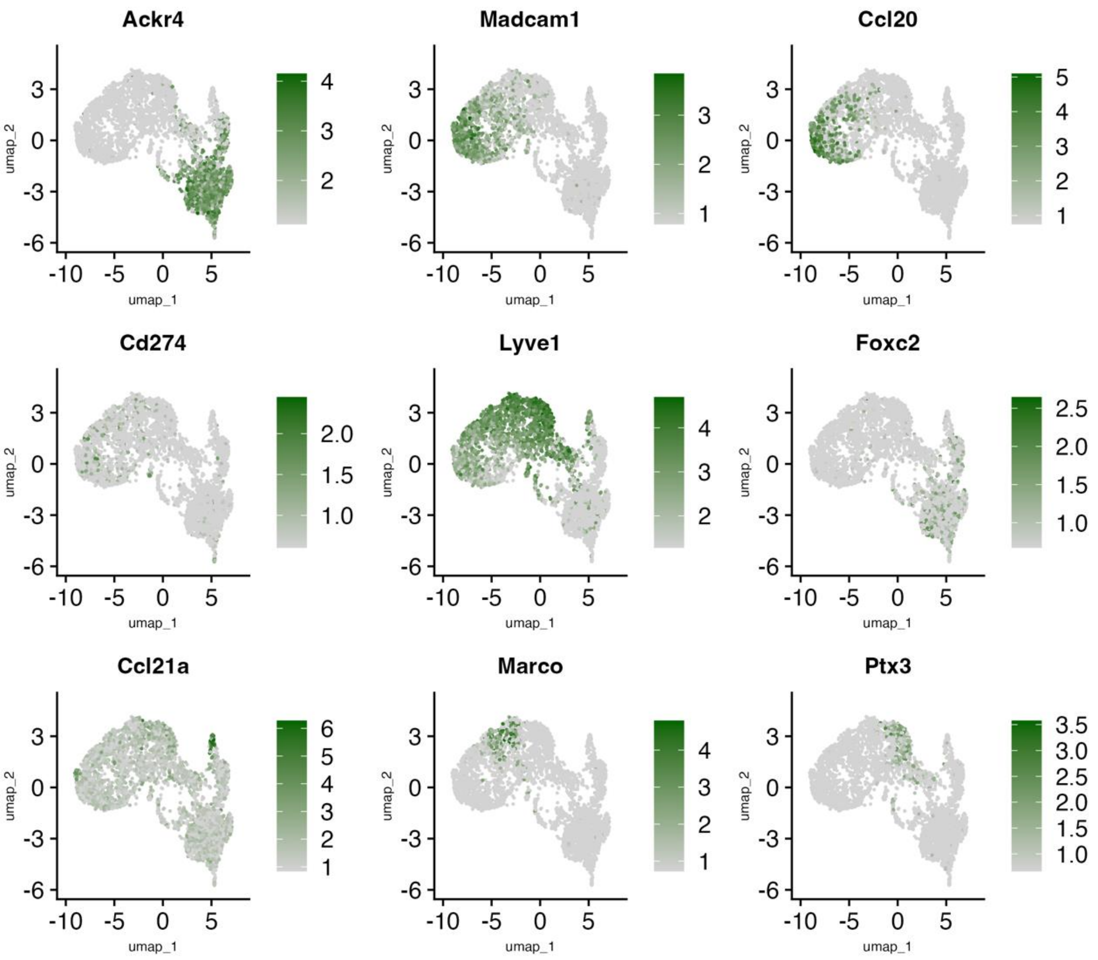


**Supplemental Figure 2.** UMAP plots of LECs from **Fig. 2A** colored by the expression of indicated marker genes.

### Supplemental Figure 3

| FRC subset | CTRL (%) | OXd4 (%) |
| --- | --- | --- |
| Ccl19lo TRC | 40.62 | 18.17 |
| Ccl19hi TRC | 11.18 | 8.93 |
| Cxcl9+ TRC | 1.29 | 1.36 |
| MRC | 14.38 | 11.06 |
| FDC | 6.19 | 6.66 |
| Cxcl13inter SC | 0.68 | 11.81 |
| dividing_cells | 0.28 | 12.60 |
| MedRC | 2.19 | 14.96 |
| Inmt+ SC | 10.26 | 8.65 |
| Cd34+ SC | 9.18 | 3.48 |
| PvC | 3.76 | 2.31 |

Changes in FRC subsets

### Supplemental Figure 4

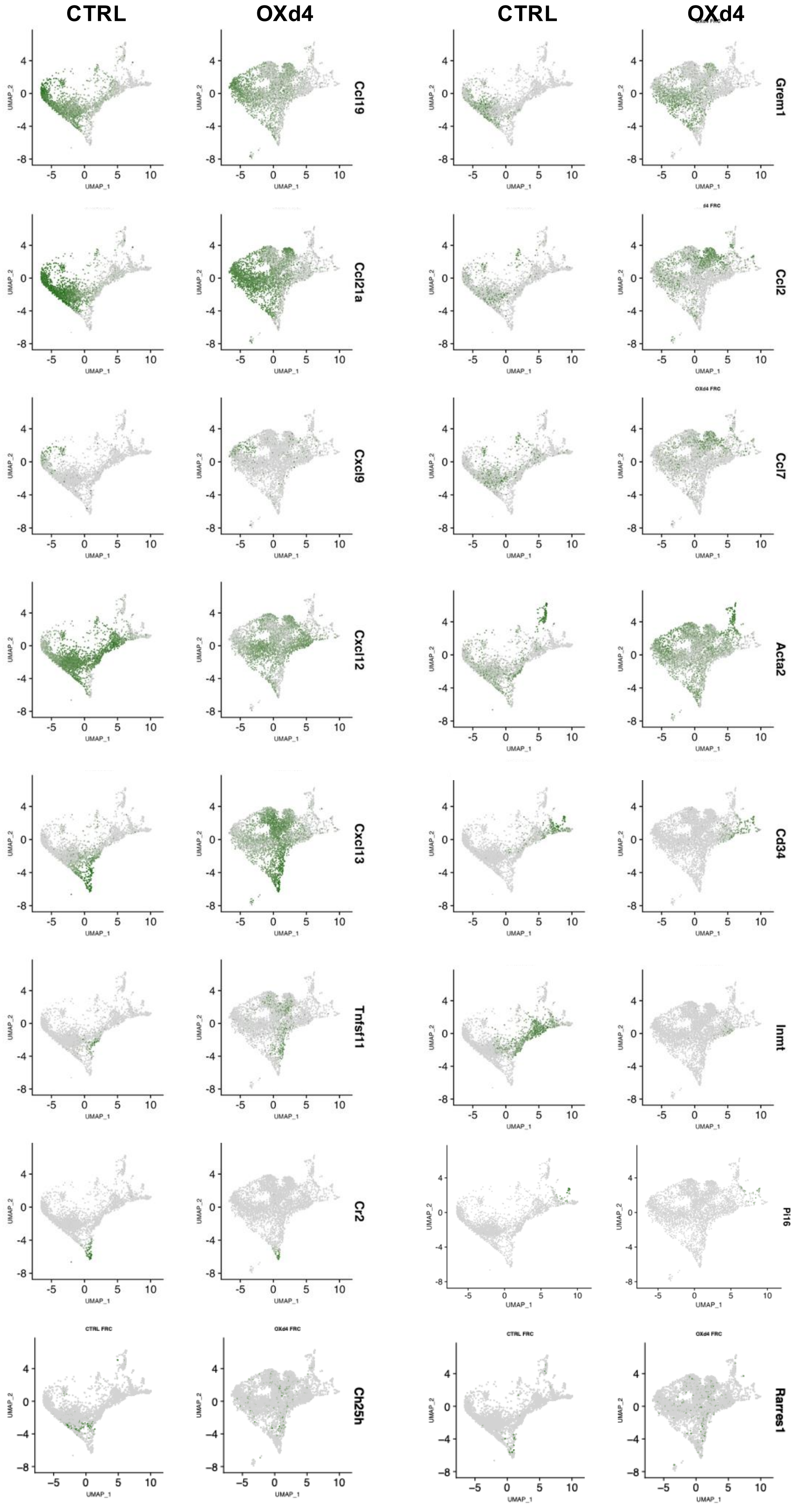


UMAP plots of FRCs from **Fig. 3B** colored by the expression of indicated marker genes.

**Supplemental Figure 5**

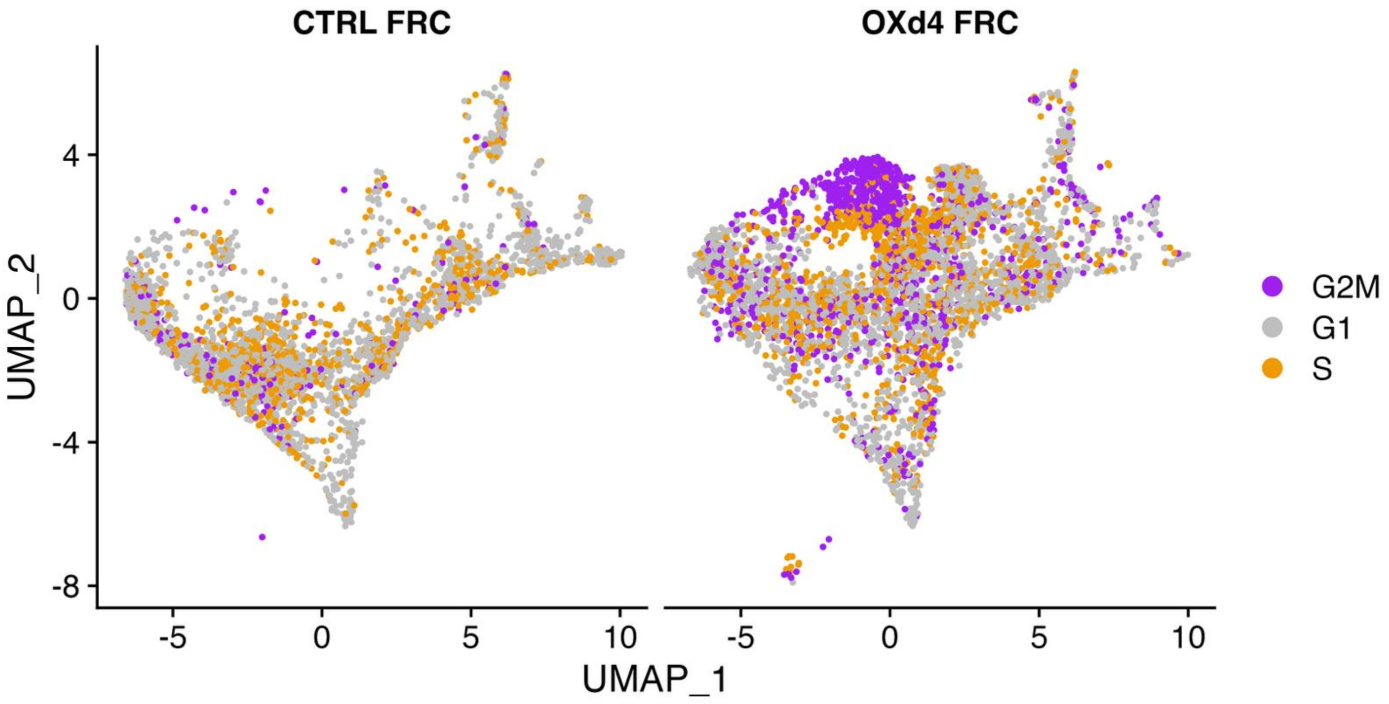


UMAP plot of FRCs from **Fig. 3B** colored by cell cycle score.

**Supplemental Figure 6**

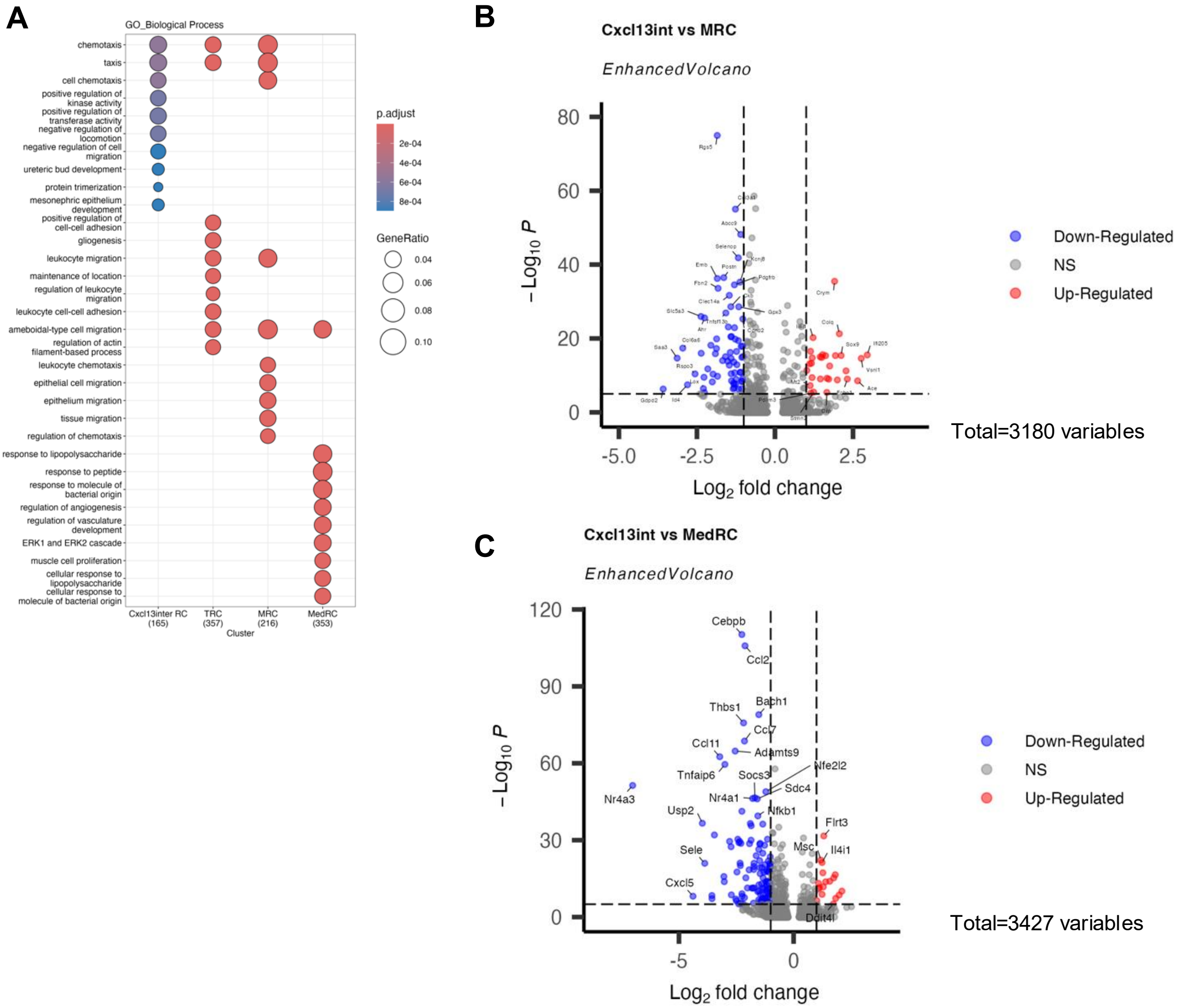


(**A**) The Dot plot shows the top 10 significant biological process pathways in Cxcl13int RC, TRC, MRC, and MedRCs in the OXd4 LNs. (**B-C**) Volcano plots show the differential gene expression in Cxcl13int RCs compared to MRCs (**B**) and MedRCs (**C**).

### Supplemental Figure 7

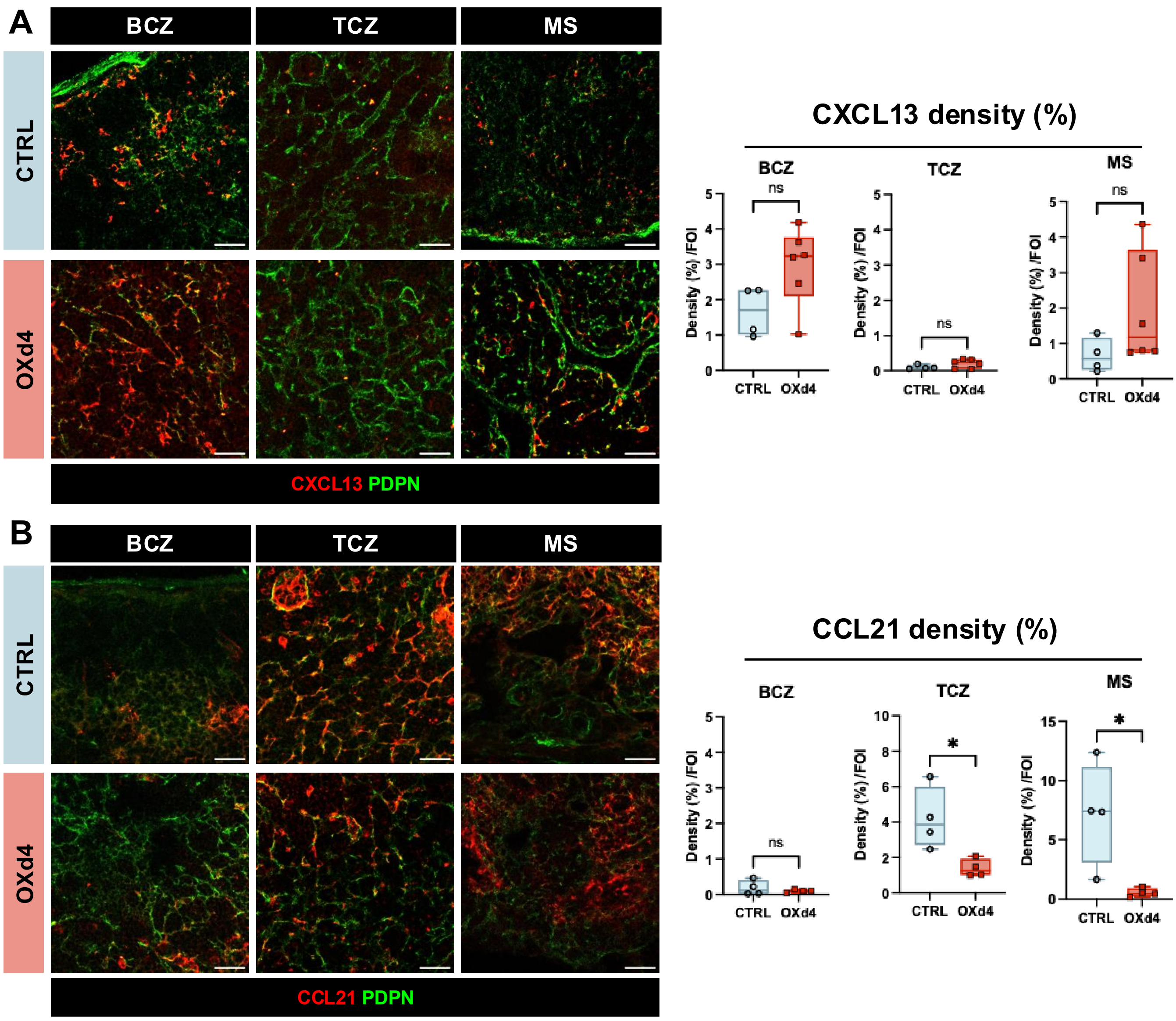


(**A**) Immunofluorescence images (left) of CXCL13 (anti-CXCL13, red) and FRCs (anti-PDPN, green) in BCZ, TCZ, and MS. Image quantification of CXCL13+ density (right). (**B**) Immunofluorescence images (left) of CCL21 (anti-CCL21, red) and FRCs (anti-PDPN, green) in BCZ, TCZ, and MS. Image quantification of CCL21 density (right). (**A-B**) Mann-Whitney test. *P < 0.05; ns, no significance.

**Supplemental Figure 8**

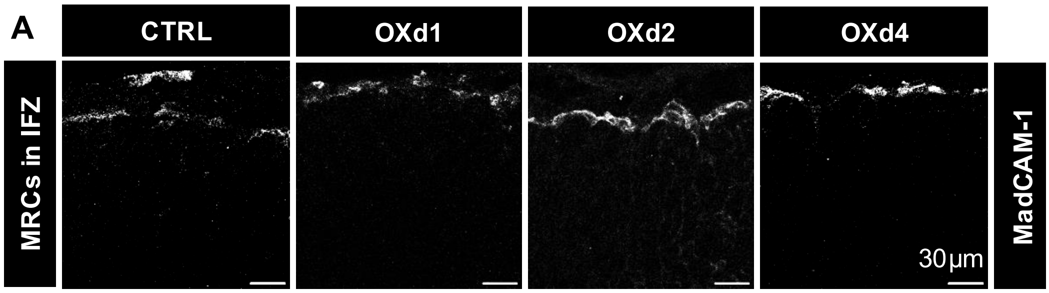


(**A**) Immunostaining of MadCAM-1 in the CTRL and OXd1-d4 LNs. N = 5-6.

**Supplemental Figure 9**

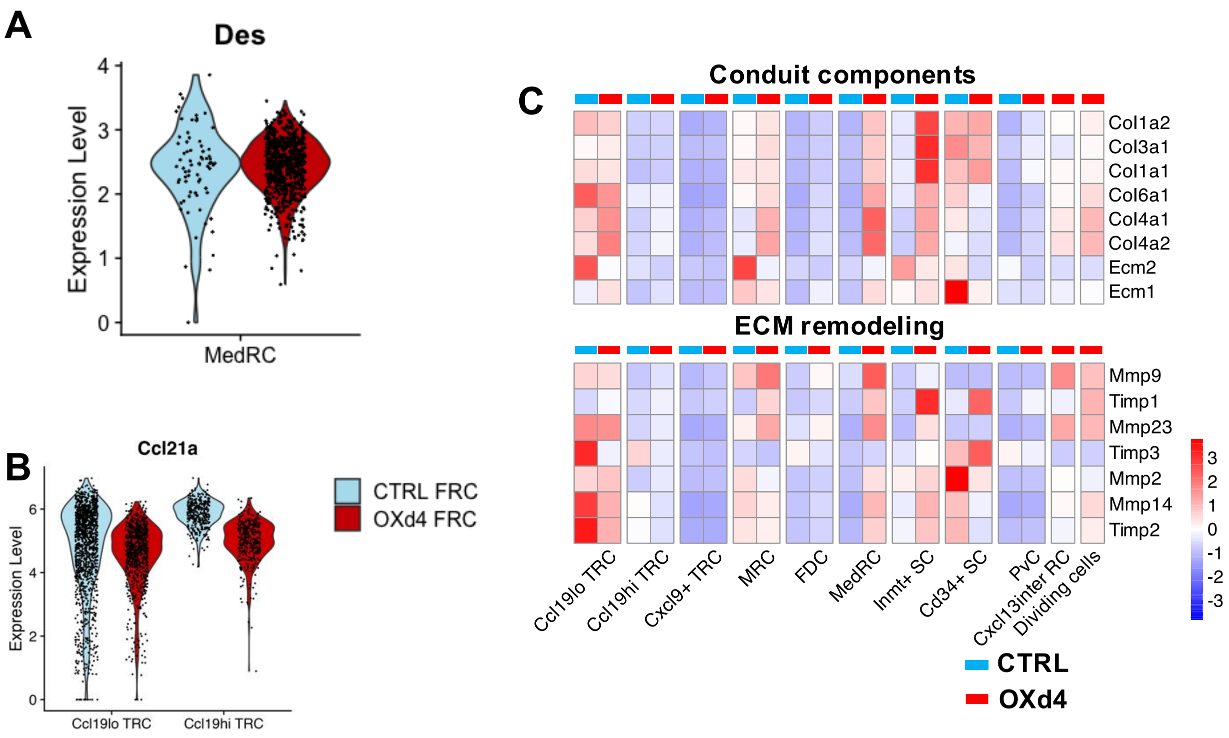


(**A**) Violin plots visualizing marker genes (*Des, Ccl21a*) expression in indicated FRC subsets. (**B**) Violin plots visualizing *Ccl21a* expression in indicated FRC subsets. (**C**) Heatmap of ECM components and ECM remodeling gene expression in FRC subsets.

**Supplemental Figure 10**

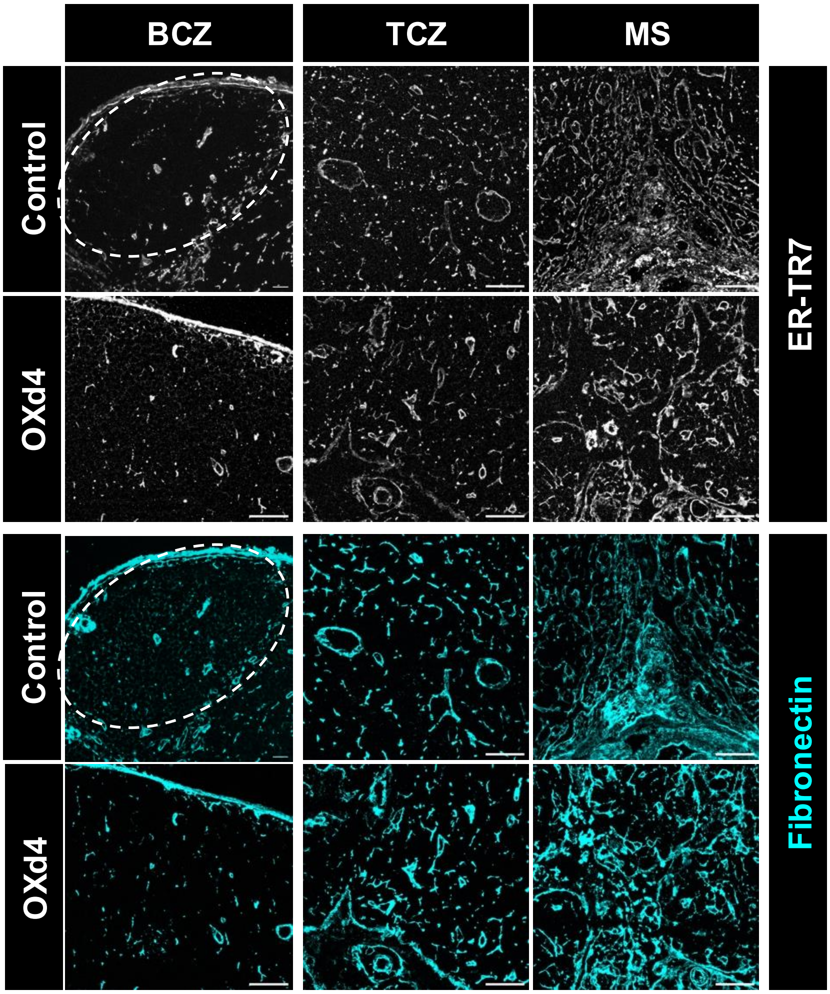


Zoom-in images of BCZ, TCZ, and MS in the CTRL and OXd4 LNs with anti-ERTR7 (grey) or anti-Fibronectin (cyan) staining.

**Supplemental Figure 11**

### CTRL FRC

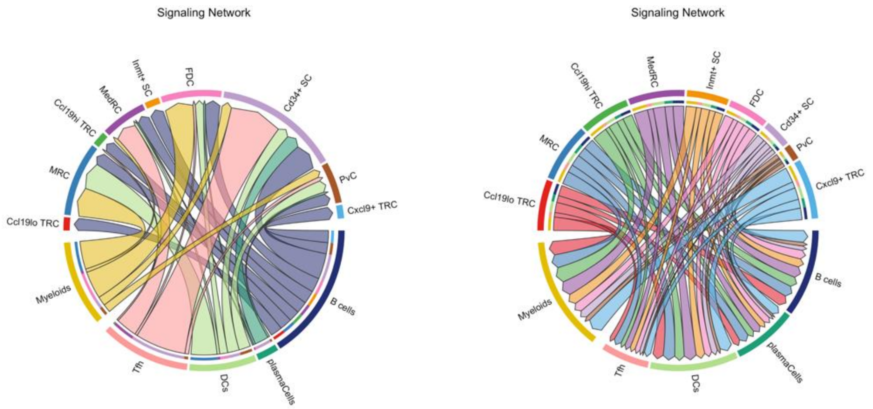


### OXd4 FRC

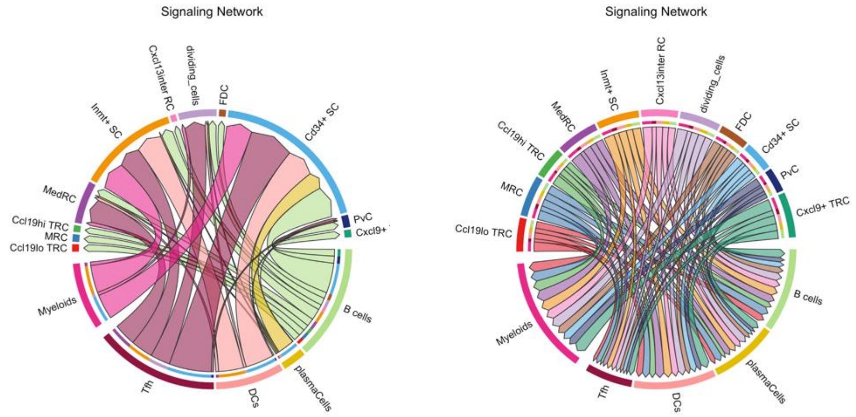


Interactome analysis in signaling pathways using CellChatDB showed the interaction between immune cells and FRCs from the CTRL (top panel) and OXd4 (bottom panel) LNs (Immune cell dataset is from M Lütge *et al.*, 2023). Left, immune cells as source and FRCs as receiver. Right, FRCs as source and immune cells as receiver.

**Supplemental Figure 12**

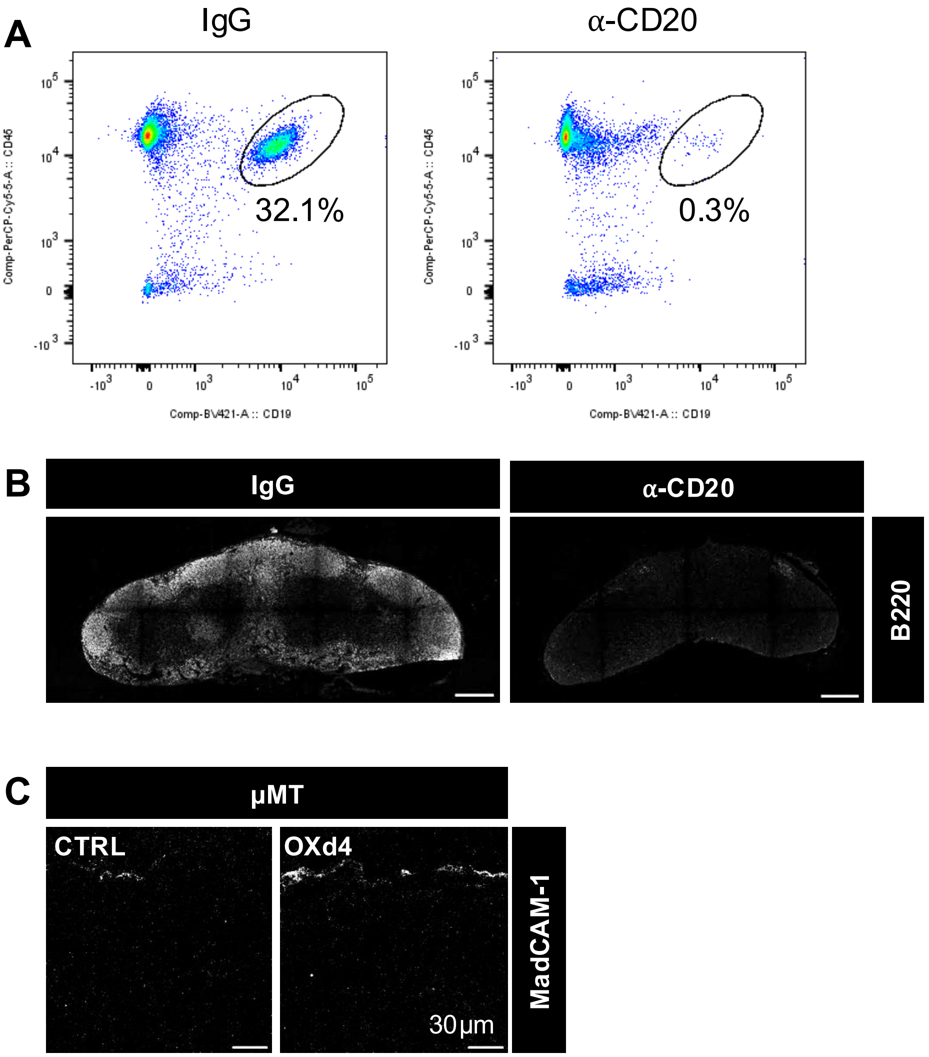


(**A-B**) Flow cytometry (**A**) and immunofluorescence staining (**B**) validated the depletion efficiency at 3 days post IgG or α-CD20 injection. (**C**) MRC staining with anti-MadCAM1 in CTRL and OXd4 LNs of µMT mice.

**Supplemental Figure 13**

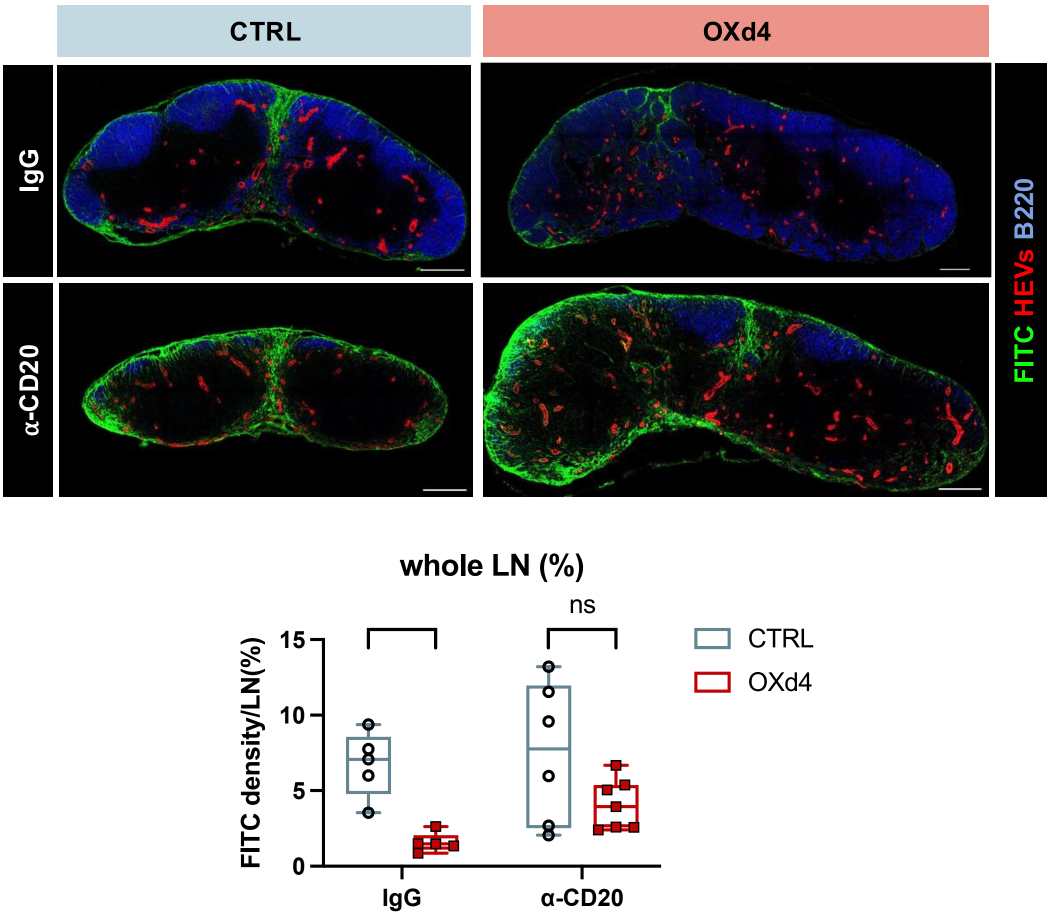


Tile scans and image quantification of total FITC drainage in CTRL and OXd4 LNs with IgG or α-CD20 treatment. N = 5-7. 2-Way ANOVA test with Šídák's multiple comparisons test. *P < 0.05; ns, no significance.

**Supplemental Figure 14**

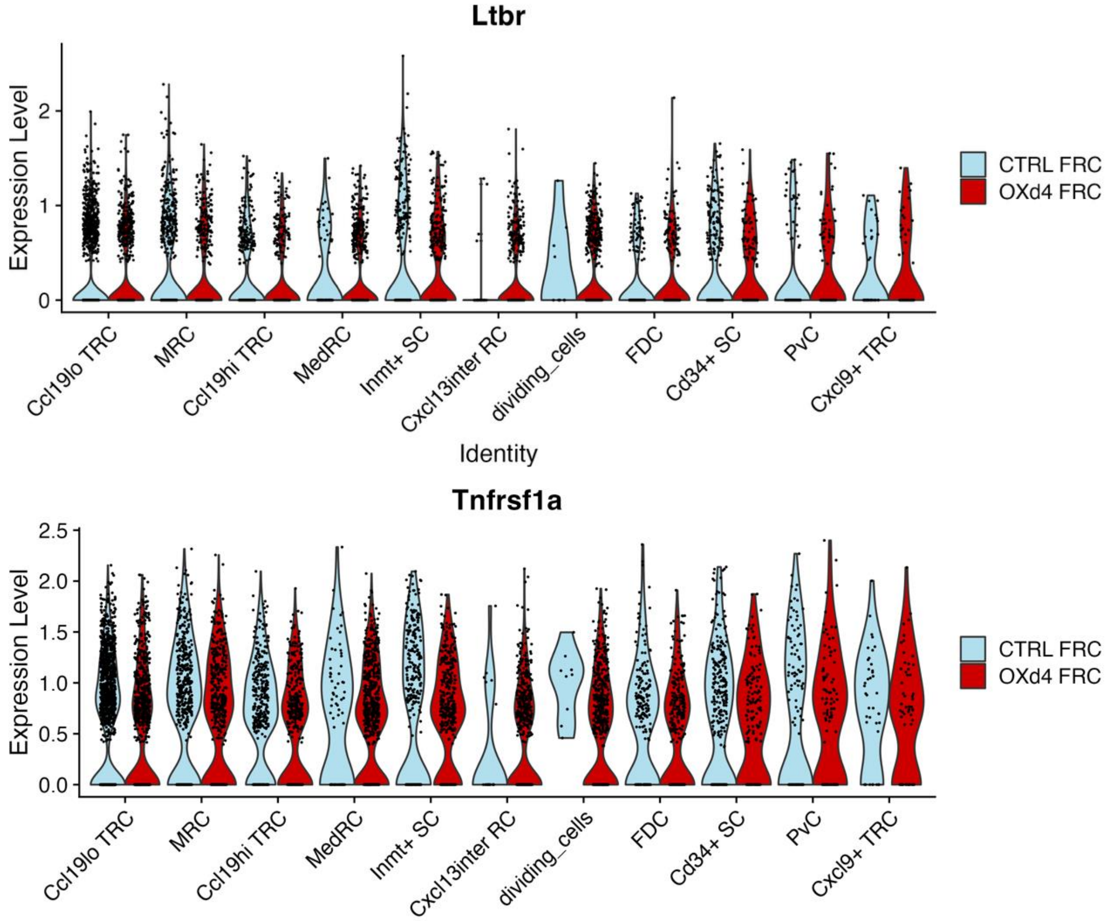


Violin plots visualizing *Ltbr* and *Tnfrsf1a* expression in FRC subsets from CTRL and OXd4 LNs.
